# Cell-type diversity and regionalized gene expression in the planarian intestine revealed by laser-capture microdissection transcriptome profiling

**DOI:** 10.1101/756924

**Authors:** David J. Forsthoefel, Nicholas I. Cejda, Umair W. Khan, Phillip A. Newmark

**Author notes:** Graduate Program in Cell and Molecular Biology, University of Wisconsin– Madison, Madison, Wisconsin, USA. Howard Hughes Medical Institute, Morgridge Institute for Research, Department of Integrative Biology, University of Wisconsin-Madison, Madison, Wisconsin, USA. Co-corresponding authors: DJF, PAN.

## Abstract

Organ regeneration requires precise coordination of new cell differentiation and remodeling of uninjured tissue to faithfully re-establish organ morphology and function. An atlas of gene expression and cell types in the uninjured state is therefore an essential pre-requisite for understanding how damage is repaired. Here, we use laser-capture microdissection (LCM) and RNA-Seq to define the transcriptome of the intestine of *Schmidtea mediterranea,* a planarian flatworm with exceptional regenerative capacity. Bioinformatic analysis of 1,844 intestine-enriched transcripts suggests extensive conservation of digestive physiology with other animals, including humans. Comparison of the intestinal transcriptome to purified absorptive intestinal cell (phagocyte) and published single-cell expression profiles confirms the identities of known intestinal cell types, and also identifies hundreds of additional transcripts with previously undetected intestinal enrichment. Furthermore, by assessing the expression patterns of 143 transcripts *in situ*, we discover unappreciated mediolateral regionalization of gene expression and cell-type diversity, especially among goblet cells. Demonstrating the utility of the intestinal transcriptome, we identify 22 intestine-enriched transcription factors, and find that several have distinct functional roles in the regeneration and maintenance of goblet cells. Furthermore, depletion of goblet cells inhibits planarian feeding and reduces viability. Altogether, our results show that LCM is a viable approach for assessing tissue-specific gene expression in planarians, and provide a new resource for further investigation of digestive tract regeneration, the physiological roles of intestinal cell types, and axial polarity.

## Introduction

Physical trauma, disease, and aging can damage internal organs. Some animals are endowed with exceptional regenerative capacity, and can repair or even completely replace damaged tissue (1–3). Regeneration requires precise spatial and temporal control over the differentiation of distinct cell types, as well as remodeling of uninjured tissue. Furthermore, individual cell types can respond uniquely to injury and play specialized signaling roles that promote regeneration (4–8). Therefore, characterization of an organ’s composition and gene expression at a cellular level in the uninjured state is an essential step in unraveling the mechanisms required for faithful re-establishment of organ morphology and physiology.

Driven by the recent application of genomic and molecular methods, the planarian flatworm *Schmidtea mediterranea* has become a powerful model in which to address the molecular and cellular underpinnings of organ regeneration (9–13). In response to nearly any type of surgical amputation injury, pluripotent stem cells called neoblasts proliferate and differentiate, regenerating brain, intestine, and other tissues lost to injury (14, 15). In addition, pre-existing tissue undergoes extensive remodeling and re-scaling through both collective migration of post-mitotic cells in undamaged tissues, as well as proportional loss of cells through apoptosis (16). These processes are coordinated to achieve re-establishment of proportion, symmetry, and function of planarian organ systems within a few weeks after injury (17).

The planarian intestine is a prominent organ whose highly branched morphology, simple cellular composition, and likely cell non-autonomous role in neoblast regulation make it a compelling model for addressing fundamental mechanisms of regeneration. In uninjured planarians, a single anterior and two posterior primary intestinal branches project into the head and tail, respectively, with secondary, tertiary, and quaternary branches extending towards lateral body margins (18, 19). Growth and regeneration of intestinal branches require considerable remodeling of pre-existing tissue (19). Remodeling is governed by axial polarity cues (16, 20), ERK and EGFR signaling pathways (21–23), cytoskeletal regulators (24), and interactions with muscle (25–28). However, the mechanisms by which post-mitotic intestinal cells sense and respond to extrinsic signals are only superficially understood.

New intestinal cells (the progeny of neoblasts) differentiate at the severed ends of injured gut branches, as well as in regions of significant remodeling, providing an intriguing example of how differentiation and remodeling must be coordinated to achieve integration of old and new tissue (19). Only three cell types comprise the intestinal epithelium: secretory goblet cells, absorptive phagocytes (29–31), and a recently identified population of basally located “outer” intestinal cells (32). Transcription factors expressed by intestinal progenitors (“gamma” neoblasts) and their progeny have been identified (24, 32–36). However, only the EGF receptor *egfr-1* (23) has been shown definitively to be required for integration of intestinal cells into gut branches, and therefore the functional requirements for differentiation of new intestinal cells are largely undefined.

The intestine may also play a niche-like role in modulating neoblast dynamics. Knockdown of several intestine-enriched transcription factors (*nkx2.2, gata4/5/6-1*) causes reduced blastema formation and/or decreased neoblast proliferation (24, 37). Similarly, knockdown of the intestine-enriched HECT E3 ubiquitin ligase *wwp1* causes disruption of intestinal integrity, reduced blastema formation, and neoblast loss (38). Conversely, knockdown of *egfr-1* causes hyperproliferation and expansion of several neoblast subclasses (23). Because these genes are also expressed by neoblasts and their progeny, careful analysis will be required to distinguish their functions in the stem cell compartment from cell non-autonomous roles in the intestine. Nonetheless, because so few extrinsic signals (39–42) controlling neoblast proliferation have been identified, further investigation of the intestine as a potential source of such cues is warranted.

Addressing these aspects of intestine regeneration and function necessitates development of approaches for purification of intestinal tissue. Previously, we developed a method for purifying intestinal phagocytes from single-cell suspensions derived from planarians fed magnetic beads, enabling characterization of gene expression by this cell type (24). More recently, single-cell profiling of whole planarians has distinguished individual intestinal cell types, as well as transitional markers for neoblasts differentiating along endodermal lineages (32, 35, 36, 43). Both approaches have advanced our understanding of intestinal biology. However, methods that (a) avoid the potentially confounding effects of feeding and dissociation on gene expression and (b) overcome the need to sequence tens of thousands of planarian cells to identify intestinal cells (only 1-3% of all planarian cells, (44)), would further enhance the experimental accessibility of the intestine.

Laser-capture microdissection (LCM) was developed as a precise method for obtaining enriched or pure cell populations from tissue samples, including archived biopsy and surgical specimens (45). Since its introduction, LCM has been used to address a vast array of basic and clinical problems requiring genome, transcriptome, or proteome analysis in specific tissues or cell types (46, 47). LCM requires sample preparation including fixation, histological sectioning, and tissue labeling or staining (48). For subsequent expression profiling, tissue processing must be optimized to maintain morphology and labeling of cells of interest, as well as RNA integrity (48–51). Excision and capture of tissue with infrared and/or ultraviolet lasers is then performed using one of several commercial LCM systems (47).

Here, we report the application of LCM for expression profiling of the planarian intestine. We first identified appropriate tissue-processing conditions for extraction of intact RNA from planarian tissue sections. Then, using RNA-Seq, bioinformatics approaches, and whole-mount in situ hybridization, we characterized the intestinal transcriptome, and discovered previously unappreciated regionalization and diversity of intestinal cell types and subtypes, especially amongst goblet cells. In addition, we identified 22 intestine-enriched transcription factors, including several required for production and/or maintenance of goblet cells, setting the stage for further studies of this cell type. The planarian intestinal transcriptome is a foundational resource for investigating numerous aspects of intestine regeneration and physiology. Furthermore, the LCM methods we introduce offer an additional, complementary strategy for assessing tissue-specific gene expression in planarians.

## Results

### Application of laser-capture microdissection to recover RNA from the planarian intestine

Successful application of laser microdissection requires identification of sample preparation conditions that balance the need to extract high quality total RNA with preservation of specimen morphology and the ability to identify tissues or cells of interest. Fixation of whole planarians requires an initial step to relax/kill animals and remove mucus, followed by fixation (52, 53). We tested three commonly used relaxation/mucus-removal treatments and two fixatives (formaldehyde and methacarn), separately and together, using short treatment times in order to minimize potential deleterious effects on RNA (Fig S1A). None of the relaxation treatments detrimentally affected RNA quality, but methacarn (a precipitating fixative) enabled much better RNA recovery than formaldehyde (a cross-linking fixative) (Fig S1A and S1B). Next, we assessed how mucus removal affected morphology and staining of cryosections taken from methacarn-fixed planarians, again using a rapid protocol to minimize RNA degradation (Fig S1C). For all three mucus-removal methods, Eosin Y alone or with Hematoxylin enabled superior demarcation of the intestine, as compared to Hematoxylin alone (Fig S1C). For preservation of morphology, magnesium relaxation was superior; NAc and HCl treatment caused tearing and detachment of intestine from the slide (Fig S1C). Finally, we assessed RNA integrity from laser-microdissected intestine and non-intestine from magnesium-treated, methacarn-fixed tissue (Fig S1D, Fig 1A and 1B), stained only with Eosin Y to further minimize processing time (Fig S1C). Although additional freezing, cryosectioning, staining, and drying steps required for LCM caused a modest decrease in RNA integrity relative to whole animals (compare Fig S1A and Fig S1D), prominent 18S/28S rRNA peaks (which co-migrate in planarians, as in some other invertebrates (54–57)) (Fig S1D) indicated that the combination of magnesium-induced relaxation, brief methacarn fixation, and rapid Eosin Y staining were suitable for LCM and extraction of RNA of sufficient quality for RNA-Seq.

**Figure 1.**
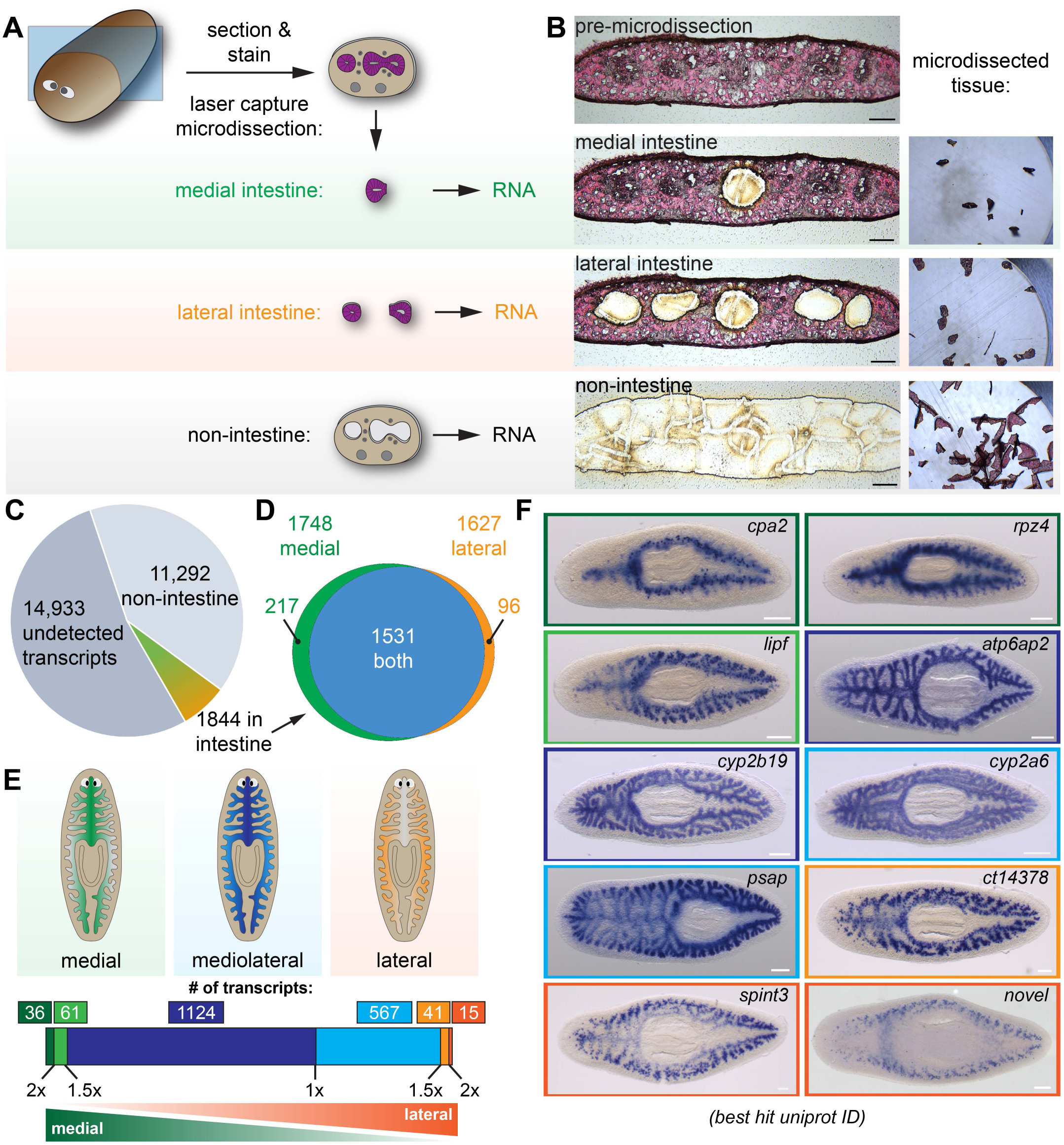
Laser-capture microdissection coupled with RNA-Seq identifies 1844 intestine-enriched transcripts. **(A)** Schematic of microdissection workflow. Planarians were fixed and cryosectioned, then sections were stained with Eosin Y. Medial intestine, lateral intestine, and non-intestine tissue were then laser captured, followed by RNA extraction and RNA-Seq. **(B)** Images of an eosin-stained section as tissue is progressively removed (left) and captured (right), yielding three samples with medial intestine, lateral intestine, and non-intestinal tissue. **(C)** Pie chart of RNA-Seq results: of 28,069 total transcripts, 13,136 were detected. Of these, 1844 were upregulated significantly in either medial or lateral intestine. **(D)** Venn diagram showing overlap of medially and laterally enriched intestinal transcripts. **(E)** Schematic of the number of transcripts with enrichment in medial or lateral intestine, expressed as a ratio of Fold Changes (FC) in each region. Dark green, FC-medial/FC-lateral > 2x. Green, FC-medial/FC-lateral = 1.5x-2x. Dark blue, FC-medial/FC-lateral = 1x-1.5x. Blue, FC-lateral/FC-medial = 1x-1.5x. Orange, FC-lateral/FC-medial = 1.5x-2x. Dark orange, FC-lateral/FC-medial > 2x. **(F)** Examples of transcripts expressed in the intestine (WISH) in medial (top), mediolateral (middle), and lateral (bottom) regions. Color outlines correspond to the color bar in panel F. Detailed gene ID information is available in Table S1 and in Results. Scale bars, 100 μm (B), 200 μm (F).

### Identification of intestine-enriched transcripts and mediolateral expression domains

Using our optimized conditions, we laser microdissected intestinal and non-intestinal tissue from four individual planarians (biological replicates) (Fig 1A, 1B and Fig S1D). For this study, we isolated tissue from the anterior of the animal (rostral to the pharynx, planarians’ centrally located feeding organ), where intestinal tissue is more abundant. We microdissected tissue from medial and lateral intestine separately, since the intestine ramifies into secondary, tertiary, and quaternary branches along the mediolateral axis, but whether gene expression varies along this axis has not been addressed systematically. We then extracted total RNA, conducted RNA-Seq, and identified transcripts that were preferentially expressed in intestinal vs. non-intestinal tissue (Fig 1C-G) (Table S1A and S1B).

Altogether, we detected 13,136 of 28,069 transcripts in the reference transcriptome (46.8% coverage) in non-intestine, medial intestine, and/or lateral intestine (Fig 1C). Of these, 1,844 were upregulated in either medial or lateral intestine, or both (Fig 1C). Specifically, in medial intestine, 1,748 transcripts were significantly upregulated (fold-change>2 and FDR-adjusted *p-*value<0.01) compared to non-intestine (Fig 1D). In lateral intestine, 1,627 transcripts were upregulated (Fig 1D). Although most (1,531/1,844) transcripts were upregulated in both medial and lateral intestine, a small subset of transcripts was significantly upregulated *only* in medial (217/1,844) or lateral (96/1,844) intestine (Fig 1D), relative to non-intestine. To further define medial and lateral transcript enrichment, we calculated a “mediolateral ratio” of the medial and lateral intestine/non-intestine fold-changes (Fig 1E and Table S1A). Although most transcripts were only modestly enriched (<1.5X fold-change enrichment) in medial (1,124) or lateral (567) intestine, a small number of transcripts was >1.5X enriched in medial (97) or lateral (56) intestine (Fig 1E and Table S1A).

We validated RNA-Seq results using whole-mount in situ hybridization (WISH) to test expression in fixed, uninjured planarians (58–60) (Fig 1F and Fig S2). 143/162 transcripts (∼88%) had detectable expression in the intestine (Fig S2, Table S1A). Most transcripts were expressed uniformly throughout the intestine (e.g., *ATPase H+-transporting accessory protein 2 (atp6ap2), cytochrome P450 2B19 (cyp2b19), cytochrome P450 2A6* (*cyp2a6),* and *prosaposin (psap)*; Fig 1F, blue borders, and Fig S2). By contrast, transcripts predicted by RNA-Seq to be most enriched (>1.5X) in medial intestine branches (e.g., *carboxypeptidase A2 (cpa2), rapunzel 4 (rpz4),* and *gastric triacylglycerol lipase (lipf),* green borders, Fig 1F) or lateral intestine branches (e.g., a *carboxypeptidase* homolog *(ct14378)*, *serine peptidase inhibitor Kunitz type 3 (spint3),* and a novel gene *(“novel”),* orange borders, Fig 1F) were indeed expressed at higher levels in these regions. Additionally, although we did not explicitly compare anterior and posterior gene expression, we also discovered anteriorly (a *C-type lectin (Zgc:171670/dd_79),* Fig S2) and posteriorly (*lysosomal acid lipase (lipa/dd_122)*, Fig S2) enriched transcripts, consistent with the influence of anteroposterior polarity cues on intestinal morphology (61–66). In summary, using LCM together with RNA-Seq identified >1800 intestine-enriched transcripts, and revealed previously unappreciated regional gene expression domains in the intestine.

### Diverse digestive physiology genes are expressed in the planarian intestine

In order to globally characterize functional classes of genes expressed in the planarian intestine, we assigned Gene Ontology (GO) Biological Process (BP) terms to planarian transcripts based on homology to human, mouse, zebrafish, *Drosophila,* and *C. elegans* genes, and then identified over-represented terms among intestine-enriched transcripts (Fig 2A, 2B, Table S1A, and Table S2A). Highly represented terms fell broadly into seven groups (Fig 2A), all of which are related to the intestine’s roles in digestion, nutrient storage and distribution, as well as innate immunity.

**Figure 2.**
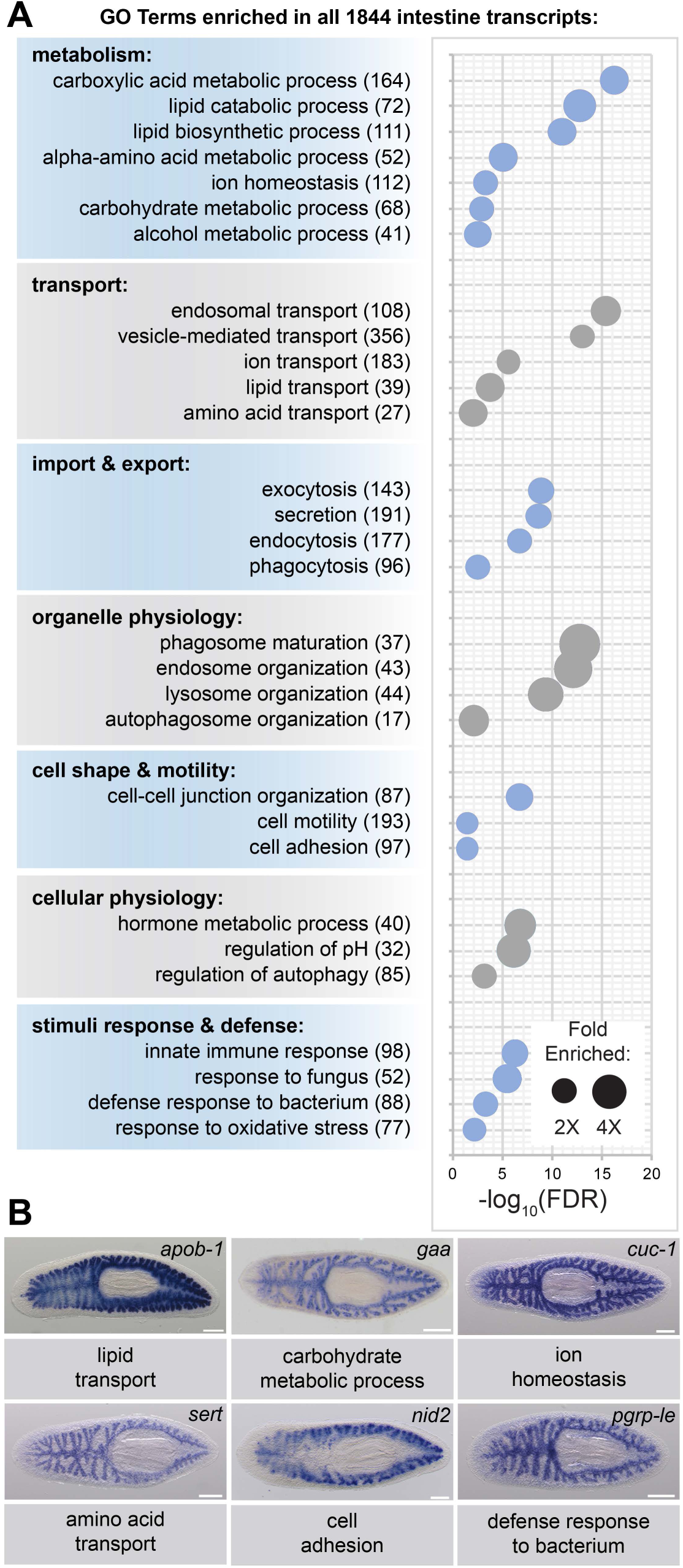
Transcripts involved in metabolism, transport, organelle physiology, transport, and stimuli responses are enriched in the intestine. **(A)** Biological Process Gene Ontology terms significantly over-represented in intestine-enriched transcripts. Bubble size indicates fold enrichment relative to all transcripts detected in laser-microdissected tissue, while position on the x-axis indicates FDR-adjusted significance. Numbers in parentheses indicate the number of intestine-enriched transcripts annotated with each term. **(B)** Examples of intestinal expression (WISH) of transcripts involved in biological processes indicated. Detailed gene ID information is available in Table S1 and in Results. Scale bars, 200 μm.

#### Metabolism

Metabolic processes were among the most highly represented: hundreds of transcripts were predicted to regulate catabolism, biosynthesis, and transport of a variety of macromolecules (e.g., lipids and carbohydrates) and small molecules (e.g., amino acids and ions). Consistent with an expected role in the initial macromolecular breakdown of ingested food, transcripts encoding secreted proteins known to regulate luminal digestion of proteins and lipids were intestine enriched, including *pancreatic carboxypeptidase A2* (a metalloexopeptidase) and three *gastric triacylglycerol lipase/lipase F* paralogs (Fig 1F and Table S1A). The intestine also is a major site of neutral lipid storage (29, 67). Transcripts related to lipid metabolism included *apolipoprotein B* paralogs (*apob-1,* Fig 2B) (regulators of lipoprotein particle biogenesis and secretion); *hormone sensitive lipase* (a regulator of diglyceride and cholesteryl ester hydrolysis); several *Niemann-Pick disease type C1/C2 protein* paralogs (intracellular cholesterol transporters); *7-dihydrocholesterol reductase* (cholesterol and steroid biosynthesis); *ceramide glucosyltransferase* (sphingolipid metabolism); and *fatty acid binding protein* (fatty acid transport), whose homologs are upregulated in the intestines of numerous animals (68–72) (Table S1A and Table S2A). Regulators of both complex and simple carbohydrate metabolism were also intestine enriched, such as the glucosidases *lysosomal alpha-glucosidase* (*gaa,* Fig 2B) and *beta-mannosidase*; *sorbitol dehydrogenase* (fructose biosynthesis); and *glycogenin-2,* which synthesizes glycogen, a complex polysaccharide found in cytoplasmic granules of phagocytes that may be an additional energy reserve (67, 73, 74) (Table S2A).

#### Transport, import, and export

Hundreds of upregulated transcripts were predicted to regulate molecular transport, vesicular trafficking, and organelle-based import and export (Fig 2A, Table S2A). Small-molecule transporters included regulators of iron transport and storage (three *ferritin* paralogs) and the intracellular copper transporter *atox1/cuc-1* (Fig 2B). Over 70 members of the solute carrier family of transmembrane transporters (e.g., *sodium-dependent serotonin transporter/sert,* Fig 2B, and Table S1) were also upregulated, reinforcing the intestine’s central role in metabolic processes, and also suggesting a potential role supporting the excretory system in maintaining extracellular solute concentration (65). Enriched regulators of vesicular trafficking included several components of clathrin-associated adaptor protein (AP) complexes, and nearly 30 Ras-related Rab GTPase proteins (Table S1, Table S2A). Other putative regulators of organelle-based import and export included homologs of *phosphatidylinositol-binding clathrin assembly protein* (*PICALM*), *epsin-1*, and several *rac* GTPase paralogs (endocytosis and phagocytosis); as well as *exocyst complex components 1, 6b, and 8, vesicle-associated membrane protein 3 (vamp3),* and *ezrin* (exocytosis and secretion). Interestingly, several transcripts encoding SNARE proteins were intestine enriched (e.g., *syntaxin-binding protein 2, extended synaptotagmin-2, synaptogyrin-1,* Table S1), as in mammalian enteroendocrine cells (75), potentially indicating an as-yet-uncharacterized role in neuroendocrine communication between intestinal cells and/or other planarian cell types.

#### Organelle and cellular physiology

Among regulators of digestion-related organelles, transcripts predicted to coordinate phagosome, endosome, and lysosome physiology were among the most highly represented. These included several *vacuolar protein sorting-associated protein (vps)* homologs required for lysosome tethering to late endosomes and autophagosomes (76), and *V-type proton ATPase 16 kDa proteolipid subunit* (*atp6v0c*), responsible for lysosome acidification (as part of the V-ATPase proton pump) and also a component of the v-ATPase-Ragulator complex that senses nutrient sufficiency/deficiency (77) (Table S1A, Table S2A). Genes regulating additional aspects of cellular physiology were also expressed in the intestine, including homologs of the autophagy-related proteins *atg3* and *atg7,* ubiquitin ligases that are required in mouse for autophagosome formation during nutrient starvation (78, 79) (Table S1A). Autophagosomes accumulate in the intestines of starved planarians (80), suggesting functional conservation of autophagy pathways. Along with *Smed-tor* (planarian mTOR), another intestine-enriched transcript (81) (Table S1A), Atg3 and Atg7 may contribute to planarians’ ability to survive extended periods of fasting (82).

#### Cell shape and motility

As in our previous study of phagocyte expression (24), here we identified many intestine-enriched genes predicted to regulate cell shape, motility, and adhesion. Expression of this class of transcripts is consistent with the intestine’s ability to remodel its branched morphology during regeneration and growth, and its role in intracellular digestion that likely requires extensive intracellular trafficking and robust maintenance of apicobasal polarity. Examples of these transcripts include numerous cytoskeletal regulators (e.g., multiple *rho-family* GTPases, *ena/VASP-like protein EVL, cofilin-1, supervillin,* and *advillin*), and regulators of cell adhesion and interactions with extracellular matrix (e.g., two *protocadherin-1-*like transcripts, *neuroglian/neural cell adhesion molecule L1-like,* two *laminin* subunits, and *nidogen-2* (Fig 2B)). Additionally, several transcripts may coordinate cell polarity and/or cell:cell junction formation, such as *lin7b,* whose orthologs stabilize mammalian cell polarity (83) and are required for cell junction targeting of the LET-23 EGF receptor during *C. elegans* vulva morphogenesis (84), and *partitioning defective 6 (pard6b),* which we previously demonstrated was required for planarian intestinal remodeling (24) (Table S1A, Table S2A).

#### Stimuli response and defense

Transcripts predicted to regulate responses to stress, microorganisms, and other stimuli were also intestine enriched (Fig 2A). Prominent among these were regulators of innate immunity, including paralogs of *peptidoglycan recognition proteins* that function in immune signaling and/or have bactericidal activity (85, 86) (*prgp-le,* Fig 2B); an ortholog of *tollip*, a modulator of responses to inflammation by innate immune cells (87) (Table S1 and Table S2); and over 30 tumor necrosis factor receptor-associated factor homologs (TRAFs), a family of adaptor proteins that is expanded in *S. mediterranea* (88) and function as effectors of receptor signaling in innate and adaptive immunity (89) (Table S1 and Table S2). *pgrp* and *traf* homologs have functional roles in planarian responses to infection with pathogenic *Pseudomonas* bacteria (90). Notably, sixty-two intestine-enriched transcripts (from this study) were previously found to be upregulated in response to shifting planarians from recirculating to static culture conditions, which causes microbiome dysbiosis (90) (Fig S3A, Table S3). Further supporting a role for the intestine in innate immunity and/or inflammatory responses, homologs of 99 *S. mediterranea* intestine-enriched transcripts were also upregulated by ingestion of pathogenic bacteria in *Dugesia japonica,* a related planarian species (91) (Fig S3B, Table S3).

### Evolutionary conservation of human digestive organ gene expression

Further illustrating the conservation of physiological roles, we found that the intestine expresses numerous homologs of transcripts that are enriched in human GI tissues (Fig 3A-F and Table S4). To make this comparison, we conducted reciprocal best homology (RBH) searches (92, 93) to identify 5,583 planarian transcripts encoding predicted open reading frames (ORFs) with high homology to ORFs in UniProt human transcripts (94), and vice versa (Fig 3A and 3B). Of these, we then identified 5,561 transcripts (including 699 gut-enriched transcripts) with human transcripts represented in the Human Protein Atlas (Fig 3B and 3C), in which transcripts whose unique or highly enriched tissue-specific expression defines 32 human tissues or organs (95). Next, we calculated the percentage of all (5,561) and gut-enriched (699) RBH transcripts that were enriched in human tissues (Fig 3D), then expressed these percentages as a fold-enrichment ratio (planarian intestine vs. non-intestine) to estimate similarity between the planarian intestine transcriptome and the transcriptomes of human tissues (Fig 3E).

**Figure 3.**
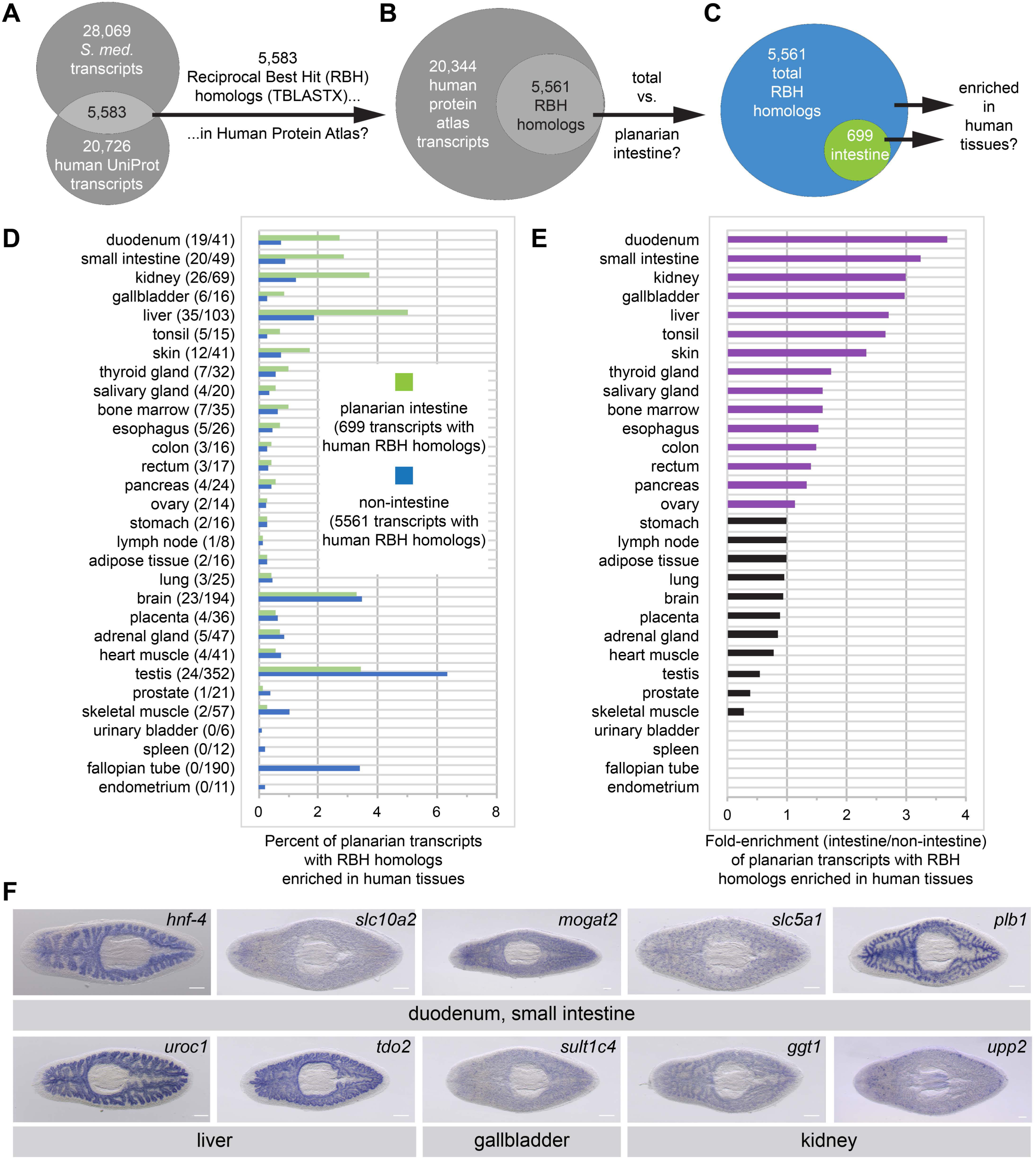
Homologs of planarian intestinal transcripts are enriched in human digestive tissues. **(A)** 5,583 *S. mediterranea* and *Homo sapiens* UniProt transcripts hit each other in reciprocal TBLASTX queries. **(B)** 5,561/5,583 human UniProt RBH homologs of planarian transcripts were present in the Human Protein Atlas. **(C)** 699/5,561 RBH homologs were enriched in the planarian intestine. **(D)** Enrichment of RBH homologs in human tissues. The first number in parentheses is the number of planarian intestine-enriched RBH homologs (of 699) and the second number in parentheses is the number of all RBH homologs (of 5,561) for each human tissue. Histogram bars represent percentage of planarian transcripts with RBH homologs enriched in human tissues. For example, 19/699 (2.72%) UniProt RBH homologs of planarian intestine-enriched transcripts were tissue-enriched, tissue-enhanced, or group-enriched in the duodenum, while only 41/5561 (0.74%) of all UniProt RBH homologs of planarian transcripts were similarly enriched. **(E)** Fold enrichment (planarian intestine/non-intestine) of RBH homologs for each human tissue. Histogram bars were calculated as a ratio of percentages in panel D. For example, 2.72% of 699 planarian intestinal RBH homologs and 0.74% of all 5,561 planarian RBH homologs were enriched in the duodenum, yielding fold enrichment of 2.72/0.74 = 3.68X. **(F)** Intestinal expression (WISH) of transcripts with RBH homologs enriched in human digestive organs: *hnf-4* (duodenum, small intestine); *slc10a2* (duodenum, kidney, small intestine); *mogat2* (colon, duodenum, liver, rectum, small intestine); *slc5a1* (duodenum, small intestine); *plb1* (small intestine); *uroc1* and *tdo2* (liver only); *sult1c14* (gallbladder only); *ggt1, upp2* (kidney only). With the exception of *Smed-hnf-4,* gene names are based on human Uniprot best hit IDs. Detailed gene ID information is available in Table S1 and in Results. Scale bars, 200 μm.

Strikingly, four of the five tissues to which the planarian intestine was most similar are involved in digestion (duodenum, small intestine, and gallbladder) or energy storage/metabolism (liver) (Fig. 3E and 3F). The intestine was also similar to other human digestive tissues (esophagus, colon, and rectum), as well as kidney, possibly suggesting a role supporting the planarian protonephridial system in filtration of extracellular solutes (Fig 3E and 3F) (65, 96, 97). Duodenum- and small intestine-enriched RBH transcripts included *hepatocyte nuclear factor 4 (hnf-4),* a transcription factor that regulates specification of endodermal tissues like intestine and liver (33), and that is expressed by a subset of planarian neoblasts likely to be intestinal precursors (32, 33, 36); the ileal (small intestine) sodium/bile transporter *slc10a2*; an ortholog of *2-acylglycerol O-acyltransferase 2 (mogat2),* an intracellular acyltransferase involved in dietary fat absorption in the small intestine (98); *sodium/glucose transporter 2 (slc5a9),* predicted to regulate glucose absorption; and a membrane-associated *phosphlipase B1* homolog (*plb1)* predicted to regulate lipid absorption (Fig. 3F). RBH homologs enriched in liver included *urocanate hydratase (uroc1)* and *tryptophan 2,3-dioxygenase* (*tdo2*), hepatic regulators of histidine and tryptophan catabolism, respectively (99, 100) (Fig 3F); and *fructose-1,6-bisphosphatase 1 (fbp1),* a regulator of gluconeogenesis and a coordinator of appetite and adiposity (101, 102) (Table S4H). In addition, orthologs of proteins involved in drug and toxic/xenobiotic compound metabolism (e.g., *ATP-binding cassette subfamily C member 2 (abcc2)* and two *cytochrome P450* family transcripts) (Tables S1A, S2A, and S4H), and lipoprotein particle biogenesis (103) (*microsomal triglyceride transfer protein,* (Table S4H) were enriched in duodenum, small intestine, and/or liver. Finally, examples of transcripts with homologs enriched in human gallbladder/kidney included *sulfotransferase 1C4 (sult1c4), gamma-glutamyltranspeptidase 1 (ggt1),* and *uridine phosphorylase 2 (upp2)* (Fig 3F). These observations reinforce the evolutionary conservation of intestinal gene expression, and indicate that physiological roles performed by multiple human digestive organs are consolidated in the planarian intestine.

### Identification of genes expressed by three distinct intestinal cell types and comparison to single-cell RNA-Seq data

Previously, we identified genes preferentially expressed by intestinal phagocytes (24). To distinguish transcripts expressed by phagocytes and other intestinal cell types, such as goblet cells (30, 74, 104), we directly compared log_2_ fold-change values for 1,317 transcripts represented in our sorted phagocyte data (24) as well as laser microdissected intestinal tissue (this study) (Fig 4A, C, and E, and Table S5). 900/1,317 transcripts were significantly upregulated in both phagocytes and laser-microdissected intestine (Fig 4A). We analyzed expression of these transcripts using WISH (Fig 4B, Table S1A, and Fig S2). As expected, these transcripts, which included previously identified intestinal markers *hnf-4* and *nkx2.2* (24, 33, 105), displayed uniform, ubiquitous expression throughout the intestine, consistent with enrichment in phagocytes (Fig 4B and Fig S2).

**Figure 4.**
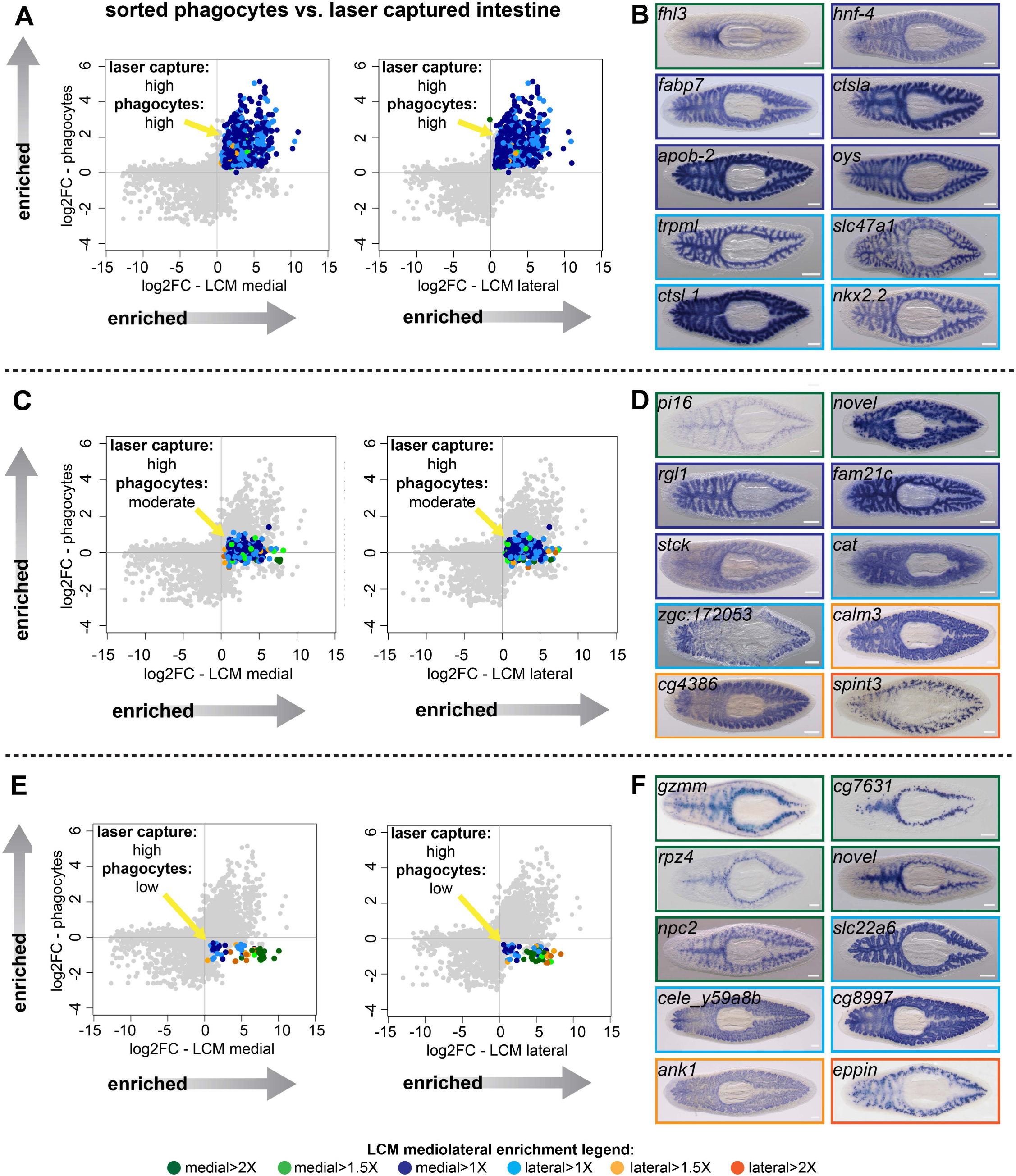
Identification of transcripts enriched in specific intestine cell types and regions. **(A)** Log_2_ fold-changes for laser-microdissected medial (left) and lateral (right) intestinal tissue (relative to non-intestinal tissue) are plotted on the x-axis, while log_2_ fold-changes for sorted/purified intestinal phagocytes (compared to all other cell types) are plotted on the y-axis. Transcripts in the upper right quadrant (colorized according to the legend in Fig 1 and at the bottom of this figure) are expressed preferentially in laser captured intestine (fold-change>2, FDR-adjusted *p* value<.01) *and* preferentially in sorted phagocytes (fold-change>2, FDR-adjusted *p* value<.05) (phagocytes: “high”). Most of these transcripts are not medially or laterally enriched in LCM transcriptomes. **(B)** Whole-mount in situ hybridizations on uninjured planarians showing examples of expression patterns for transcripts in (A). Borders are colorized according to the mediolateral legend in Fig 1 and at the bottom of the figure. Expression patterns are mostly uniform and ubiquitous in the intestine, consistent with phagocyte-specific expression, with the exception of *fhl3* (top left), which is medially enriched. **(C)** Plots as in (A), but with colorized transcripts expressed preferentially in laser-captured intestine, but not significantly up- or down-regulated in sorted phagocytes (FDR-adjusted *p* value>.05) (phagocytes: “moderate”). Some of these transcripts are medially or laterally enriched in LCM transcriptomes. **(D)** Examples of gene expression for transcripts in (C). A variety of intestine expression patterns is observed. **(E)** Plots as in (A), with transcripts in the lower right quadrant enriched in laser microdissected intestine, but significantly downregulated (fold-change <2, FDR-adjusted *p* value<.05) in sorted phagocytes relative to non-phagocytes (phagocytes: “low”). Many of these transcripts are enriched in medial or lateral LCM transcriptomes. **(F)** Examples of gene expression patterns for transcripts in (E). A majority of these transcripts are enriched in goblet or basal cells, sometimes in medial or lateral subpopulations. Detailed gene ID information is available in Table S1 and in Results. Scale bars, 200 μm.

358 intestine-enriched transcripts were not significantly up- or down-regulated in phagocytes (Fig 4C, Fig S2, and Table S5). WISH analysis suggested these transcripts are enriched in multiple intestinal cell types (Fig 4D). Some transcripts in this group were expressed ubiquitously throughout the intestine (*ral guanine nucleotide dissociation stimulator-like 1 (rgl1)* and *family with sequence similarity 21 member C (fam21c),* a homolog of *WASH complex subunit 2,* Fig 4D), suggesting expression in phagocytes, possibly in addition to other cell types. However, others were expressed in a distinct subset of less abundant intestinal cells (*peptidase inhibitor 16 (pi16)* and *serine peptidase inhibitor, Kunitz type 3 (spint3)*, Fig 4D). These transcripts are enriched in goblet cells, since their WISH expression pattern is highly similar to labeling of this subpopulation by lectins (106), antibodies (53, 64, 107), and other recently identified transcripts (32, 43, 64). A third set of transcripts was enriched in basal regions of the intestine (*zgc:172053,* a homolog of human C-type lectin *collectin-10,* and *calmodulin-3 (calm3)*, Fig 4D). This pattern resembles that of a planarian *gli*-family transcription factor (108) and several solute carrier-family transporters (65), and indicates expression by “outer intestinal cells” (which we refer to as “basal cells” because of their proximity to the basal region of phagocytes) that were also recently identified in a large-scale, single-cell sequencing effort (32).

Finally, 59 intestine-enriched transcripts were significantly downregulated in phagocytes (Fig 4E, Fig S2, and Table S5). As expected, we never observed uniform, phagocyte-like expression patterns for these transcripts (Fig 4F and Fig S2). Rather, transcripts in this group were enriched only in goblet cells (e.g., *epididymal secretory protein E1/Niemann-Pick disease type C2 protein (npc2)*) or basal cells (e.g., *solute carrier family 22 member 6 (slc22a6)*). Furthermore, some goblet-cell-specific transcripts also appeared to be either medially (e.g., *metalloendopeptidase (cg7631)*) or laterally (e.g., *eppin*) enriched (Fig 4F and Fig S2), suggesting possible specialization of this cell type along the mediolateral axis.

Overall, validation by WISH identified 78 transcripts with a ubiquitous intestinal expression pattern; nearly all of these were upregulated in our sorted phagocyte data (Fig S2 and Fig S4A). By contrast, most of the 25 validated goblet-cell-enriched transcripts (Fig S4B) and 14 basal-cell-enriched transcripts (Fig S4C) were not upregulated in sorted phagocytes.

We further validated our results by comparing transcript enrichment in enterocytes/phagocytes, goblet cells, and outer intestinal cells/basal cells reported in a recent single-cell RNA-Seq (scRNA-Seq) analysis of planarian cells (32). Phagocyte- and goblet-specific transcripts mapped to similar quadrants in our phagocyte vs. laser-captured intestine plots (S4D and Fig S4E). Most outer intestinal cell-specific transcripts were downregulated in our phagocyte data set, but a few were upregulated (Fig S4F), suggesting that basal cells and phagocytes may share greater transcriptome similarity than phagocytes and goblet cells. A second scRNA-Seq study (43) also identified phagocyte- and goblet-cell-specific transcripts. Again, these mapped to similar quadrants in our phagocyte vs. laser-captured intestine plots (Fig S4G-H). Furthermore, we identified 809 intestine-enriched transcripts that single-cell studies did not find to be enriched in the intestine (or for some, any planarian cell type) (Fig S4I, Table S1A). Using WISH, we validated intestine enrichment for 23/28 of these mRNAs (Table S1A), including some with a uniform/phagocyte-like expression pattern (e.g., *sodium-dependent serotonin transporter (SerT)* (Fig 2B); *sodium/calcium exchanger 1 (slc8a1)* (Fig S4I); and *PITH-domain containing protein CG6153 (cg6153)* (Fig S4I)), and also goblet-enriched transcripts (e.g., *carboxypeptidase A2 (cpa2)* (Fig 1F) and *kallikrein 13 (klk13)* (Fig S4I). We also note that over 1000 intestine-enriched transcripts in scRNA-Seq studies were not included in our LCM-generated transcriptome (Fig S4I). However, the vast majority (>80%, not shown) of these were enriched in multiple cell types (32, 43) (Table S1A), suggesting considerable expression in non-intestinal tissue, consistent with our data.

Taken together, our results confirm the existence of three major intestinal cell types that have unique, but overlapping, gene expression profiles (32, 43). These include phagocytes and goblet cells, as well as a third population of cells that resides in basal regions of intestinal branches. In addition, the detection of transcripts exclusively enriched in laser-captured intestine suggests that expression profiling of laser-captured bulk tissue is significantly more sensitive than current single-cell profiling approaches, and may be a preferable method for assessing tissue-specific gene expression when single-cell resolution is not required.

### Multiple cell types and subtypes reside in the planarian intestine

To further characterize intestinal cell types and subtypes, we used fluorescence in situ hybridization (FISH) to investigate co-expression of intestine-enriched transcripts. First, we verified the existence of three distinct cell types, using highly expressed phagocyte, goblet, and basal-specific markers (Fig 5A-C). Expression of the most phagocyte-enriched transcript, *cathepsin La (ctsla)*, was ubiquitous throughout intestinal branches, but did not overlap with *npc2*, a goblet-cell-enriched mRNA (Fig 5A), or with *slc22a6,* a basally enriched transcript (Fig 5B). A second phagocyte-enriched transcript, *cathepsin L1 (ctsl.1),* was co-expressed with *ctsla,* but not with *npc2* or *slc22a6* (Fig S5A). A third phagocyte-enriched mRNA, *apolipoprotein b-2 (apob-2),* was expressed in more basal regions of *ctsla+* phagocytes, but again, *apob-2* expression was clearly distinct from *npc2+* goblet cells and *slc22a6+* basal cells (Fig S5A). Goblet cell-enriched *npc2* was expressed by cells with minimal *slc22a6* expression (Fig 5C), further reinforcing that *npc2+* goblet cells are distinct from *ctsla+* phagocytes as well as *slc22a6+* basal cells. We also found that basal cells were distinct from numerous visceral muscle fibers that surround intestinal branches, occupying basal regions around digestive cells (109, 110), consistent with another study (28). Basal-specific *slc22a6* mRNA did not colocalize with either a muscle-specific mRNA (*troponin I 4 (tni-4),* Fig 5D) (5), or with labeling by an antibody that recognizes a muscle-specific antigen (mAb 6G10, Fig 5E) (53). Thus, *slc22a6+* cells represent a novel cell type in the intestine that is distinct from visceral muscles, phagocytes, and goblet cells, and that has, to our knowledge, not been described by numerous previous histological and ultrastructural studies. Our data independently confirm the identification of this cell type in a recent single-cell sequencing effort (32).

**Figure 5.**
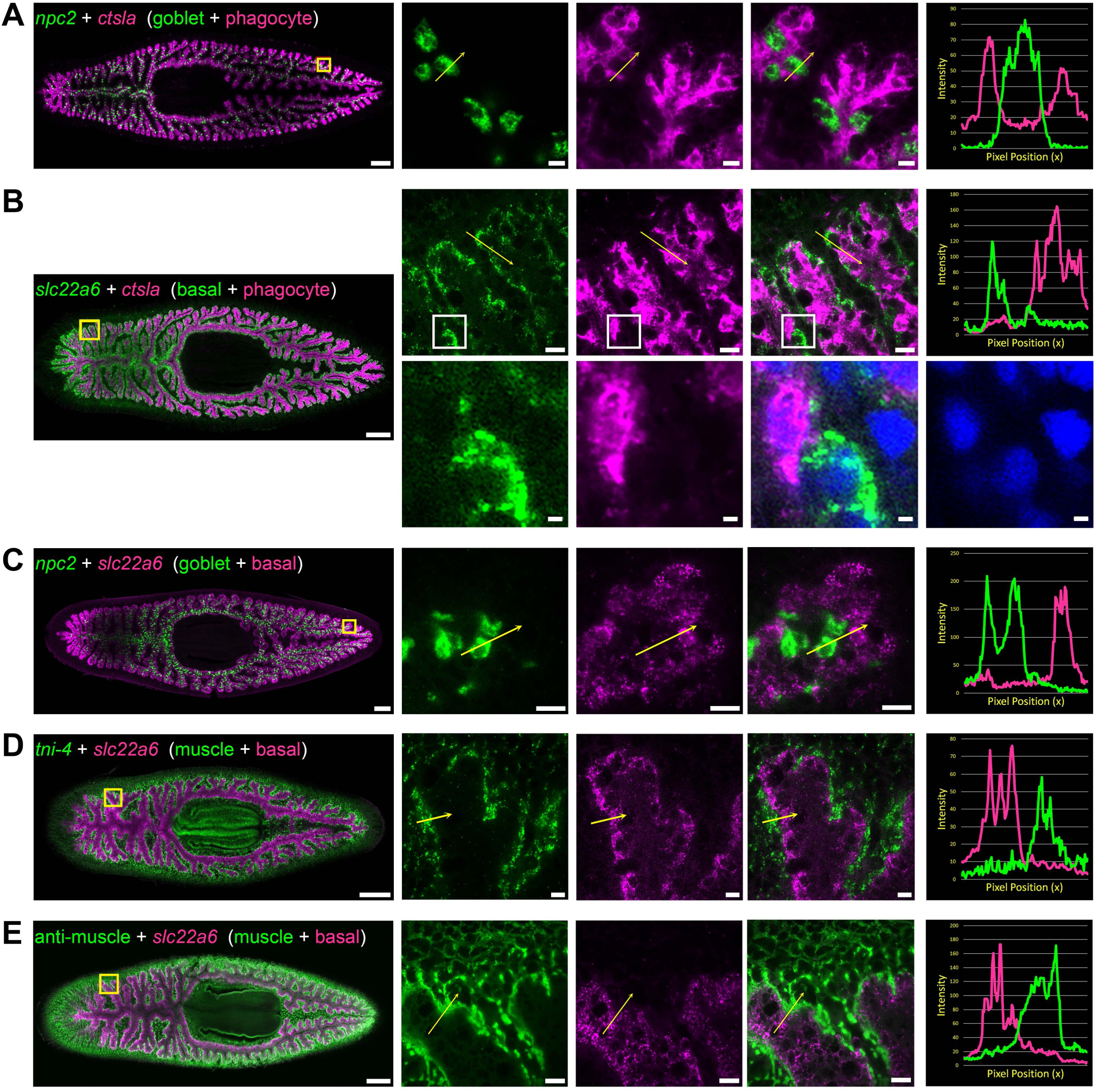
Double fluorescence in situ hybridization reveals three major cell types in the planarian intestine. **(A)** Confocal images of *npc2* (green) and *ctsla* (magenta) in situ hybridization. Left to right, whole animal (yellow box indicates magnified region in right panels), zoomed area in green, magenta, and merge (yellow arrow indicates profile line in right-most panel), and a graph showing pixel intensity in each color from tail to head of the yellow profile arrow. *ctsla* is the top phagocyte-specific gene in the phagocyte microarray dataset, while *npc2* is enriched in goblet cells. **(B)** *slc22a6* (green) and *ctsla* (magenta). *slc22a6* mRNA is restricted to the basal region of the intestine, and shows minimal overlap with the phagocyte marker *ctsla*. The white box represents the cropped region shown below with DAPI labeling nuclei, indicating that these riboprobes label distinct cells. **(C)** *npc2* (green) and *slc22a6* (magenta). *npc2* is enriched in goblet cells while *slc22a6* is enriched in basal cells, with minimal overlapping signal. **(D)** *tni-4* (green) and *slc22a6* (magenta). *tni-4* is expressed by visceral muscles, while *slc22a6* is found in basal cells. **(E)** Anti-muscle antibody (6G10, green) and *slc22a6* (magenta). Detailed gene ID information is available in Table S1 and in Results. Scale bars, whole animals 200 μm; magnified images, 10 µm, magnified crop of basal cell (B), 2 µm.

Using additional markers, we also discovered previously unappreciated heterogeneity in gene expression amongst both goblet and basal cells. For example, *metalloendopeptidase (cg7631)* was expressed by a subpopulation of goblet cells restricted to medial regions of the intestine, mainly localized to primary branches (Fig 6A and 6C). By contrast, *spint3-*expressing cells were found laterally, within secondary, tertiary, and quaternary branches (Fig 6B and 6C). Only rarely did cells in these medial and lateral domains co-mingle, or co-express both markers; such occurrences were restricted to the mediolateral boundaries between primary and secondary branches (Fig 6C). We also identified subpopulations of basal cells. For example, *zgc:172053* (*collectin-10-like,* see also Fig 4D) was expressed by some, but not all basal cells, especially in lateral regions of the intestine (Fig 6D). Similarly, *si:dkey-241* (another C-type lectin) was enriched in a subset of laterally enriched basal cells, but not phagocytes or goblet cells (Fig S5B). These data reveal subpopulations of goblet and basal cells in distinct intestinal domains, suggesting potentially specialized roles.

**Figure 6.**
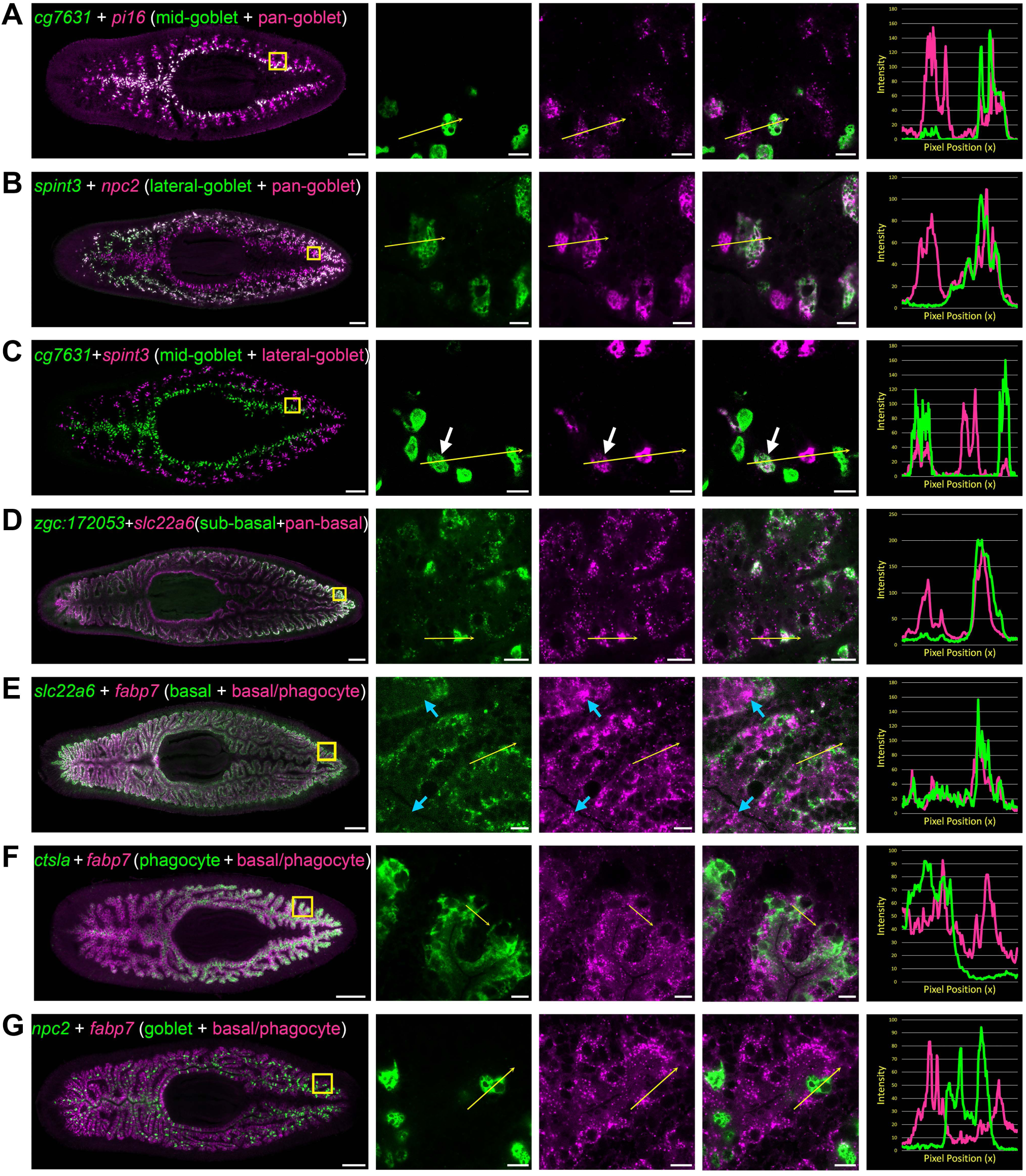
Transcripts expressed by intestinal subpopulations and multiple cell types. **(A)** Confocal images of *cg7631* (green) and *pi16* (magenta) in situ hybridization. Left to right, whole animal (yellow box indicates magnified region in right panels), zoomed area in green, magenta, and merge (yellow arrow indicates profile line in right-most panel), and a graph showing pixel intensity in each color from tail to head of the yellow profile arrow. *cg7631* is enriched in medial goblet cells, while *pi16* is found in all goblet cells. **(B)** *spint3* (green) and *npc2* (magenta). *spint3* is enriched in the lateral goblet cell population, while *npc2* is expressed by all goblet cells. **(C)** *cg7631* (green) and *spint3* (magenta). *cg7631* is enriched in medial goblet cells; *spint3* is enriched in lateral goblet cells. Only rarely do these two markers label the same cell, indicated with a white arrow. **(D)** *zgc:172053* (green) and *slc22a6* (magenta). *zgc:172053* is enriched in a subset of basal cells, while *slc22a6* is more ubiquitously enriched in most basal cells. **(E)** *slc22a6* (green) and *fabp7* (magenta). *slc22a6* is a basally enriched gene, while *fabp7* is expressed by both basal cells and more apical cells (phagocytes). Blue arrows indicate apical gene expression where *slc22a6* is absent. **(F)** *ctsla* (green) and *fabp7* (magenta). *ctsla* expression is enriched in phagocytes, while *fabp7* is found in both phagocytes and basal cells. **(G)** *npc2* (green) and *fabp7* (magenta). *npc2* is enriched in goblet cells, and overlaps minimally with *fabp7* in phagocytes and basal cells. Detailed gene ID information is available in Table S1 and in Results. Scale bars, whole animals 200 μm; magnified images, 10 µm.

We also identified transcripts expressed by multiple cell types. For example, *fatty acid binding protein 7* (*fabp7)* mRNA was expressed by both basal cells (Fig 6E) and phagocytes (Fig 6F), but negligibly in goblet cells (Fig 6G). By contrast, *lysosomal alpha-glucosidase (gaa)* was expressed at high levels in goblet cells, at low levels in phagocytes, and at negligible levels in basal cells (Fig S5C). *oysgedart (oys),* a lysophospholipid acyltransferase, was enriched in phagocytes, and at low levels in basal cells, but not in goblet cells (Fig S5D). These results, together with bioinformatic comparisons (Fig 4 and Fig S4), demonstrate that unique digestive cells co-express many transcripts in different combinations and levels, illustrating the complexity of gene expression even in a tissue with relatively few cell types.

### Intestine-enriched transcription factors regulate goblet cell differentiation and maintenance

Definition of the intestinal transcriptome enables identification of genes required for regeneration and functions of distinct intestinal cell types. Here, to initiate this effort, we focused on goblet cells, which expressed the majority of medially and laterally enriched transcripts we identified (Table S1A, Fig S2). Planarian goblet cells (44, 111) (also called “Minotian gland cells” (112), “granular club” cells (30), “sphere cells” (31), or “pyriform cells” (113)) have been proposed to secrete digestive enzymes into the intestinal lumen (104), or store protein reserves (29). Ultrastructurally, planarian goblet cells possess numerous large proteinaceous granules and abundant rough endoplasmic reticulum (30, 104, 113), resembling mammalian goblet cells that produce a protective mucous barrier and mount innate immune responses (114–118). However, although numerous markers and reagents have been identified that label planarian goblet cells (32, 43, 53, 64, 106, 107), to our knowledge, genes required for goblet cell differentiation, maintenance, or physiological roles have not been reported.

Reasoning that transcription factors (TFs) would regulate goblet cell generation and/or maintenance, we focused on transcripts encoding 22 intestine-enriched TFs, only 10 of which were previously known to be enriched in intestinal cells (Table S6A). Using FISH, we validated expression of all but one of these TFs in the intestine (Fig S6). In a dsRNA-mediated RNA interference screen to specifically assess goblet cell regeneration, we found that knockdown of three TFs dramatically reduced expression of a goblet cell marker in regenerating head, trunk, and tail fragments (Table S6A and Fig 7). Knockdown of *mediator of RNA polymerase II transcription subunit 21 (med 21)* reduced goblet cells and blastema formation (Table S6A), but also caused severe disruption of intestinal integrity in our previous study (24). This suggested a non-goblet-cell-specific role, and we did not investigate *med21* further. Knockdown of a second transcription factor, *gli-1* (a transducer of hedgehog signaling (108, 119)), caused failure of goblet cells to regenerate at the midline of the new anterior branch in amputated tail fragments regenerating a new head (Table S6A; Fig 7A and 7B; Fig S7A). Regeneration of goblet cells was also reduced in new tail branches of *gli-1(RNAi)* head fragments (Fig S7B) and in anterior and posterior branches in *gli-1(RNAi)* trunk fragments (Fig S7C). These effects on goblet cells were specific, since phagocytes (Fig 7A, Fig S7A-C) and basal cells (Fig 7B) regenerated normally. This was true even in some fragments with smaller posterior blastemas characteristic of reduced hedgehog signaling (Fig S7C) (108, 119, 120).

**Figure 7.**
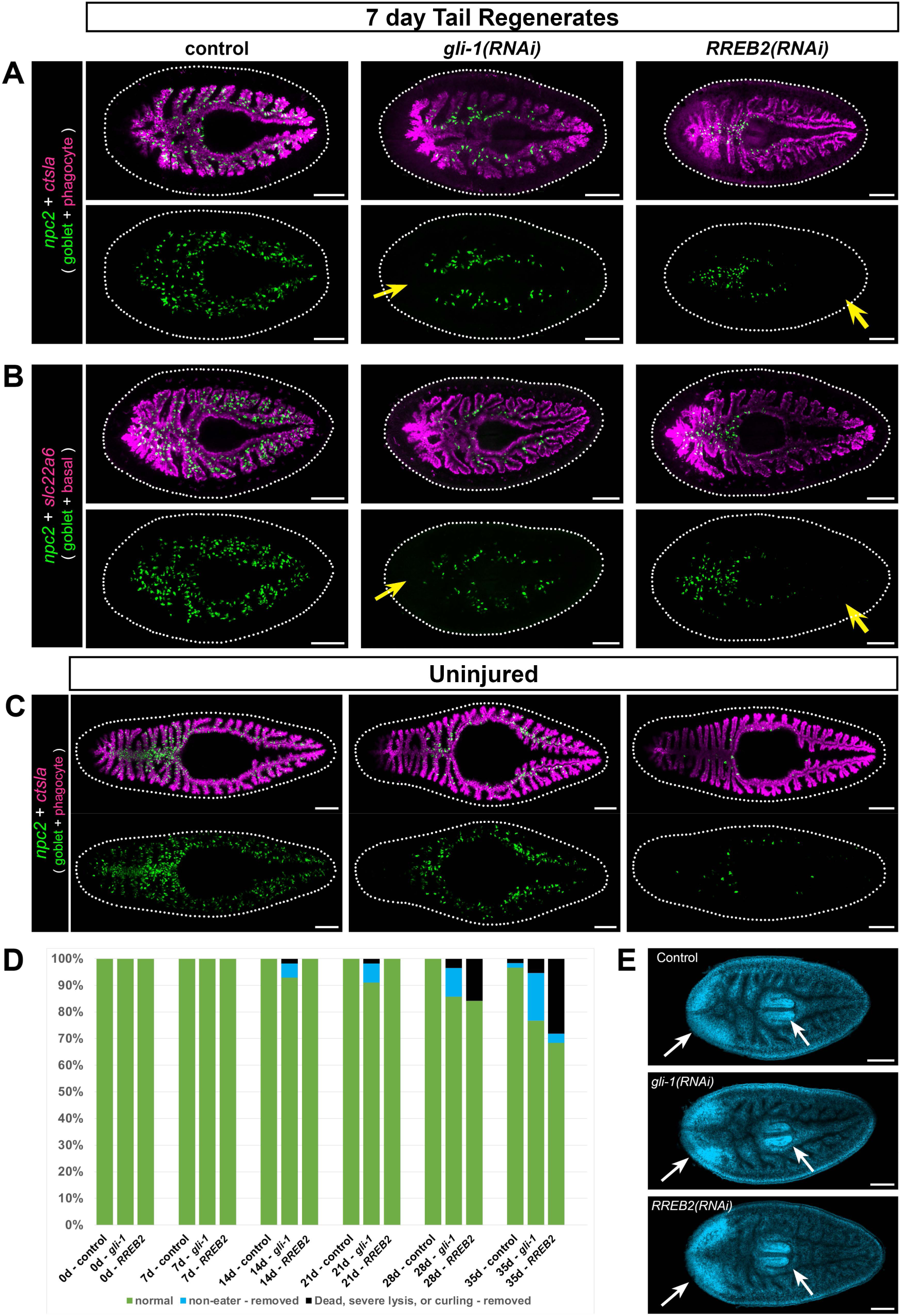
*gli-1* and *RREB2* regulate goblet cell numbers. **(A)** In 7-day tail regenerates, *gli-1* knockdown dramatically reduces goblet cells (*npc2+*) at the midline in regenerating intestine (yellow arrow), while *RREB2* knockdown reduces goblet cells in old tissue (yellow arrow). Phagocytes (*ctsla+*) appear normal in all conditions. **(B)** Basal cells (*slc22a+*) are unaffected in *gli-1(RNAi)* and *RREB2(RNAi)* regenerates, while goblet cells are reduced similar to A. **(C)** In uninjured animals, *gli-1* RNAi causes moderate goblet cell loss, while *RREB2* RNAi results in severe goblet cell loss. **(D)** Phenotypes in *gli-1(RNAi)* and *RREB2(RNAi)* planarians during 6 dsRNA feedings (once per week). Animals refuse food and undergo lysis and death with increasing frequency over the RNAi time course. Total sample size was n ≥ 55 for each condition; data were pooled from 3 independent biological replicates of n ≥18 each. **(E)** DAPI labeling of tail regenerates shown in B. White arrows indicate normal regeneration of new brain and pharynx. Animals in A, B, and E were fed dsRNA 8 times (twice per week), starved 7 days, amputated, then fixed 7 days later. Animals in C and D were fed dsRNA 6 times (once per week), starved 7 days, then fixed for FISH. Detailed gene ID information is available in Table S1 and in Results. Scale bars, 200 μm.

By contrast to the *gli-1* phenotype, goblet cells arose normally in regenerating intestine upon knockdown of a third TF, *ras-responsive element binding protein 2 (RREB2),* including anterior intestinal branches in tail fragments (Fig 7A and 7B; Fig S7A), posterior branches in head fragments (Fig S7B), and both anterior and posterior branches in trunk fragments (Fig S7C). However, in pre-existing intestinal regions, goblet cell numbers were dramatically reduced. These included the posterior of tail fragments (Fig 7A and 7B; Fig S7A), the anterior of head fragments (Fig S7B), and central regions of trunk fragments (Fig S7C). As with *gli-1,* phagocytes and basal cells were unaffected (Fig 7A and 7B; Fig S7A-C). Thus, the *RREB2* phenotype represents a striking “mirror image” of the phenotypes in *gli-1(RNAi)* animals. Together, these results suggest that *gli-1* regulates neoblast fate specification and/or differentiation of neoblast progeny into goblet cells, while *RREB2* may control maintenance or survival of goblet cells after they initially differentiate.

In uninjured animals, both *gli-1* and *RREB2* also reduced goblet cell numbers. Knockdown of *gli-1* reduced goblet cell numbers in uninjured animals, especially in anterior, posterior, and lateral intestine branches (Fig 7C), but many goblet cells remained in medial, primary intestinal branches. In *RREB2(RNAi)* animals, goblet cells were more dramatically reduced in all regions of the intestine (Fig 7C). Although these phenotypes were broadly consistent with observations in regenerates, they also suggested that *gli-1* and *RREB2* might be required for differentiation and/or maintenance of distinct mediolateral goblet cell subpopulations in uninjured animals. Indeed, we found (using additional goblet cell markers) that almost no lateral goblet cells remained in *gli-1(RNAi)* uninjured animals, while again, medial goblet cells were much less affected (Fig S8A). By contrast, in *RREB2(RNAi)* uninjured animals, reduction of both medial and lateral goblet cell numbers was pronounced, but nonetheless some lateral goblet cells remained (Fig S8A). These differential effects on mediolateral subpopulations were not observed in regenerates. For example, both subpopulations failed to differentiate in the primary anterior intestinal branch in *gli-1(RNAi)* tail regenerates (Fig S8B), and both subpopulations were severely reduced in pre-existing, posterior branches of *RREB2(RNAi)* tail regenerates (Fig S8B).

Taken together, these results suggest that, in uninjured animals undergoing normal homeostatic growth and renewal, *gli-1* is primarily required for differentiation of new lateral intestinal cells. Alternatively, as goblet cells fail to renew, remaining goblet cells might somehow migrate to more medial intestinal branches, or survival of goblet cells in primary branches could be prolonged. Conversely, *RREB2* seems to be required for maintenance of both medial and lateral goblet cells, although lateral cell numbers are slightly less affected by *RREB2* knockdown, compared to *gli-1*. Finally, in regenerates, *gli-1* and *RREB2* knockdown affects medial and lateral goblet cells similarly (but in regenerating vs. pre-existing intestine). These complex observations suggest that mechanisms regulating goblet cell differentiation and maintenance may differ during homeostasis and regeneration, or that compensatory mechanisms capable of sustaining goblet cell production/survival in uninjured *gli-1(RNAi)* and *RREB2(RNAi)* animals are insufficient to meet the increased demand for new tissue during regeneration.

Despite their specific effects on goblet cells, neither *gli-1* nor *RREB2* mRNAs are specifically enriched in this cell type. In fact, *gli-1,* which was previously shown to be expressed by intestine-associated cells (108, 121), was most highly enriched in phagocytes, basal cells, and intestine-associated muscle cells (visceral muscle), but was also expressed at lower levels in some goblet cells (Fig S9A). *RREB2* was enriched in goblet cells and basal cells, but expression was also observed in some phagocytes and intestine-associate muscle (Fig S9B). Intriguingly, we found that knockdown of two other TFs, *48 related 1 (fer1)* (also called *pancreas transcription factor 1 subunit alpha* or *PTF1A* (*32*)), and *LIM homeobox 2 (lhx2b),* also modestly reduced goblet cells in pre-existing branches (Table S6A, Fig S10A-C). mRNAs encoding these TFs are enriched in basal regions of the intestine (Fig S6). In addition, single-cell transcriptome data suggest *PTF1A* expression is elevated in differentiating neoblast progeny in the basal cell lineage (32), and *PTF1A* also reduces basal intestinal cell numbers (32). Thus, although our data support a role for *gli-1* and *RREB2* in goblet cells and/or their precursors, they also raise the possibility that basal cells (and possibly phagocytes or muscle cells) may non-autonomously influence goblet cell differentiation and/or survival.

### Goblet cell reduction compromises feeding behavior and viability

We asked whether goblet cell depletion affected planarian viability, behavior, or regeneration. Over a 6-week dsRNA feeding regimen, some uninjured *gli-1(RNAi)* and *RREB2(RNAi)* planarians refused food and failed to eat after 4-5 weeks (Fig 7D). In both *gli-1(RNAi)* and *RREB2(RNAi)* animals, some animals eventually lysed, curled, or died (Fig S11A), suggesting that goblet cells are required for viability. Additionally, we also observed a modest decrease in animal size (Fig S11B-C) in *RREB2(RNAi)* planarians relative to controls. Interestingly, we only observed feeding failure and other phenotypes in animals fed the dsRNA/liver mix 1x/week (every 7 days) for 6 weeks; animals that were fed 2x/week (every 3-4 days), but still 6 times, did not refuse to eat, lyse, curl, or die (not shown). This suggests that goblet cells may regulate hunger (or other aspects of digestive physiology) primarily in starved animals. Next, to assess regeneration, we amputated planarians fed 2x/week for 8 feedings, in order to eliminate the influence of feeding failure on possible regeneration phenotypes. In both *gli-1(RNAi)* and *RREB2(RNAi)* regenerates, goblet cell depletion was robust (Fig 7A and 7B), but neither gene was required more generally for regeneration (with the infrequent exception of reduced posterior blastemas in *gli-1(RNAi)* regenerates, mentioned above), as the brain (Fig 7E), pharynx (Fig 7E), other intestinal cell types (Fig 7A and 7B), and new intestinal branches (Fig 7A and 7B) all regenerated without noticeable defects. Together, these results suggest that goblet cells are dispensable for regeneration, and that their primary role may be to regulate appetite or other aspects of intestinal physiology that are critical for viability.

### Goblet cells likely have multiple functions

The intestinal transcriptome (Table S1A) includes numerous goblet-cell-enriched transcripts that provide insights into the potential physiological roles of goblet cells (Fig S12A and S12B). First, in support of previous suggestions that goblet cells promote lumenal digestion (111, 112, 122), we identified several goblet-enriched transcripts predicted to encode secreted regulators of protein catabolism (*pancreatic carboxypeptidase A2*) and triglyceride catabolism (*lipase F/gastric triacylglycerol lipase)* (Fig S12A). Second, we also identified three goblet-enriched *kallikreins* (*klk*) (Fig S12A and Fig S2), secreted proteases whose mammalian homologs produce vasoactive plasma kinin, but also hydrolyze extracellular matrix molecules, growth factors, hormone proteins, and antimicrobial peptides in numerous tissues (123). Kallikreins are expressed by goblet cells in rat, cat, and mouse intestines (124, 125), and have been implicated in inflammatory bowel disease and gastrointestinal cancers (126, 127), suggesting additional conservation of planarian goblet cell physiology. Third, goblet cells express *peptidoglycan recognition protein (pgrp-1b)* (Fig S12A), whose vertebrate and invertebrate homologs modulate innate immune signaling and play direct bactericidal roles (85, 86), suggesting goblet cells may coordinate immune responses and/or regulate microbiome composition. Consistent with this idea, a second planarian paralog, *pgrp-1e* (Fig 2B and Fig S2), is upregulated in the intestine (but not restricted to goblet cells) in response to *Pseudomonas* infection (90). Fourth, a homolog of *prohormone convertase (pcsk1*) is enriched in goblet cells, as well as peripharyngeal cells surrounding the pharynx (Fig S12B). Prohormone convertases (PCs) cleave neuropeptide and peptide hormone preproteins to generate bioactive peptides. In both vertebrates and invertebrates, PCs function in neurons, but also in endocrine cell types such as pancreatic islet cells and digestive tract enteroendocrine cells, where they regulate glucose levels, energy homeostasis, and appetite (125, 128, 129). Although another planarian *pcsk* paralog, *Smed-pc2,* processes neuropeptides required for germline development (130), to our knowledge, no intestine-enriched prohormones have been reported. Nonetheless, *pcsk1* expression suggests goblet cells may have an enteroendocrine-like role, consistent with our observation (Fig 7D) that goblet cell reduction perturbs feeding behavior. Finally, because mucin production is a defining feature of goblet cells in vertebrates (117) we also identified three planarian homologs of genes encoding gel-forming mucins, including MUC2, an abundant intestinal mucin (Fig S12C) (131, 132). Somewhat surprisingly, these were expressed by peripharyngeal cells that send projections into the pharynx (52) (Fig S12C), but not goblet cells. Thus, although mucin proteins may be delivered to the intestinal lumen through the pharynx, negligible goblet cell expression of *mucin* transcripts is a notable difference between planarians and vertebrates.

Intriguingly, many transcripts with the greatest medial or lateral enrichment (e.g., >2X in medial vs. lateral tissue or vice versa) in the intestine were expressed by goblet cells (Table S6B-C, Fig 4C-F, Fig S4B, Fig S2), suggesting possible functional specialization. Such transcripts included the digestive enzymes, proteases, and pattern recognition receptor just described, as well as transcripts encoding predicted extracellular matrix-associated proteins (e.g., *nidogen-2* and *eppin*) (Fig 2B, Fig 4F, Fig S2, and Table S6B-C). To attempt to determine the functional significance of mediolateral goblet cell subpopulations, we used RNAi to assess the roles of 16 of the most medial and 8 of the most lateral goblet-enriched transcripts (Table S6B-C). However, we did not observe failure to feed, decreased viability, or defects in blastema formation, even after 8 weeks of knockdown (feeding 1x/week) (Table S6B-C). This might be due to functional redundancy, since multiple lipases, carboxypeptidases, and kallikriens are expressed by goblet cells. Alternatively, other intestinal cell types might play overlapping roles with respect to some functions. For example, analysis of biological process GO term enrichment among all 1221 medial and 623 lateral transcripts – without regard to cell type specificity – revealed numerous putative regulators of innate immunity and macromolecular catabolism among medial transcripts, and of extracellular matrix organization among lateral transcripts (Fig S13A-C). Lastly, some goblet cell functions may be required only when planarians are challenged by stresses like bacterial infection or extended starvation, possibilities that will require further investigation.

## Discussion

We have developed methods for applying laser microdissection to planarian tissue, which we used to define gene expression in the intestine. The intestine expresses genes involved in metabolism, nutrient storage and transport, innate immunity, and other physiological roles, demonstrating considerable functional homology with digestive systems of other animals, including humans. Comparison of gene expression in microdissected tissue to that of intestinal phagocytes (previously isolated by sorting) enabled identification of transcripts enriched in two other cell types, goblet cells and basal cells. We also discovered previously unappreciated intestinal cell-type diversity, especially amongst goblet cells, which reside in distinct medial and lateral intestinal domains. Identification of medially and laterally enriched transcripts suggests an additional paradigm for addressing axial influences on organ regeneration, as well as evolution of patterning mechanisms influencing the regionalization of bilaterian digestive systems (133, 134). Finally, we identified intestine-enriched transcription factors that play distinct roles in goblet cell differentiation and maintenance, and found that depletion of goblet cells reduces planarians’ willingness to feed and viability, with only negligible effects on regeneration of other intestinal cell types or non-intestinal tissues.

Characterization of the planarian intestinal transcriptome provides a framework for several next steps. First, a detailed description of intestinal cell types will facilitate further studies of digestive cell types. Here, we identified two transcription factors, *gli-1* and *RREB2,* whose RNAi phenotypes suggest that distinct mechanisms govern differentiation and maintenance of goblet cells. We are unaware of studies in other organisms that have uncovered direct roles for either *gli-1* or *RREB2* in goblet cells. However, in mice, modulation of Sonic Hedgehog (an upstream activator of Gli-family TFs) levels affects goblet cell numbers (135, 136), and RREB1 cooperatively regulates expression of the peptide hormone secretin in enteroendocrine cells (137). By contrast, in the *Drosophila* midgut, Hedgehog (Hh) signaling promotes intestinal stem cell proliferation (138), while the RREB-1 ortholog *hindsight/pebbled* suppresses midgut intestinal stem cell proliferation and is required for differentiation of absorptive enterocytes (139). In planarians, hedgehog (Hh) regulates anteroposterior polarity, neoblast proliferation, and neurogenesis (108, 120, 121, 140); our data suggest a possible additional role for Hh in intestinal morphogenesis. Similarly, although a second planarian RREB paralog, *RREBP1,* is partially required for eye regeneration and may promote differentiation (141), implication of *RREB2* in goblet cell maintenance/survival suggests that Ras-mediated signaling could also influence cellular composition in the intestine (although direct links to Ras remain to be established). Thus, although characterization of the precise roles of *gli-1* and *RREB2* will be required, our results suggest broad similarity between planarians and other animals with respect to evolutionarily conserved signaling pathways that govern cell dynamics in the digestive tract. Furthermore, although we focused on goblet cells, previous studies suggest that several genes (*gata4/5/6-1, hnf4, egfr-1, PTF1A*) expressed by cycling neoblast subpopulations (23, 33, 34, 36) may be required for differentiation of multiple intestinal cell types (23, 32, 37, 142). Thus, enumeration of intestinal cell types and subtypes here, along with endoderm-specific progenitors in several recent single-cell analyses (32, 36, 43), provides a rich list of candidate regulators for further elucidation of differentiation, maintenance, and functions of planarian digestive cells and their progenitors.

Second, functional analysis of intestine-enriched transcripts will help to resolve the intestine’s role in regulation of the stem cell microenvironment. Knockdowns of the intestine-enriched TFs *nkx2.2* or *gata4/5/6-1* result in reduced proliferation and blastema production (19, 37). In addition, a subset of *tetraspanin group-specific gene-1 (tgs-1)*-positive neoblasts likely to be pluripotent neoblasts resides near intestinal branches (36), further hinting at a niche-like role for the intestine. Numerous intestine-enriched transcripts encode regulators of metabolite processing and transport, as well as putatively secreted proteins, suggesting multiple possible cell non-autonomous influences on neoblast dynamics. Because gut-enriched TFs like *nkx2.2* and *gata4/5/6-1* are also expressed by neoblasts and other cell types (32), LCM also provides an efficient approach for clarifying which candidate stem cell regulators are dependent on these or other TFs for their intestine-specific expression.

Third, LCM provides a complementary method for assessing injury-induced gene expression changes in the intestine and other planarian tissues that may overcome the shortcomings of other available methods. Previously, we developed a method for purification of intestinal phagocytes from planarians that ingested magnetic beads (24, 52). Laser microdissection may be preferable, because it enables detection of transcripts expressed by other intestinal cell types (not just phagocytes), and also eliminates potentially confounding gene expression changes caused by feeding. Similarly, although single-cell sequencing (SCS) approaches in planarians have driven significant advances in our understanding of planarian cell types and their responses to injury (32, 34, 36, 43, 143, 144), LCM may provide a more direct and efficient way to assess gene regulation in tissues with rarer cell types. For example, because intestinal cells comprise only 1-3% of total planarian cells (44), without prior enrichment SCS would potentially require analysis of tens of thousands of cells to reliably detect gene expression changes in the intestine. In addition, although chemistry and computational approaches for SCS are improving rapidly (145, 146), LCM may enable more sensitive detection of low-copy transcripts or subtle fold changes in bulk tissue, and also bypass “noise” caused by dissociation or other SCS-related technical artefacts.

Regeneration of digestive organs is not well understood. Development of a robust method for isolating intestinal tissue, and characterization of the intestinal transcriptome, will facilitate mechanistic studies in planarians. The methods and resources presented here will also support comparative analyses, complementing current and future efforts to understand digestive tract regeneration in platyhelminths, sea cucumbers, annelids, ascidians, amphibians, and mammals (147–156). In addition, studies in planarians and other regeneration models are likely to generate new insights into cellular processes (e.g., proliferation, differentiation, metabolism, and stress responses) whose dysregulation underlies human gastrointestinal pathologies associated with aging and disease.

## Materials and Methods

### Ethics Statement

No vertebrate organisms were used in this study.

### Planarian maintenance and care

Asexual *Schmidtea mediterranea* (clonal line CIW4) (157) were maintained in 0.5 g/L Instant Ocean salts with 0.0167 g/L sodium bicarbonate dissolved in Type I water (158), and fed with beef liver paste. For all experiments, planarians were starved seven days prior to fixation. For LCM, planarians were 6-9 mm in length. For WISH and FISH, planarians were 2-4 mm in length. All animals were randomly selected from large (300-500 animals) pools, with the exception that animals with blastemas (e.g., those that had recently fissioned) were excluded.

### Optimization of planarian fixation for RNA extraction and histology

Planarians were relaxed in 0.66 M MgCl_2_ or treated with 7.5% N-acetyl-L-cysteine or 2% HCl (ice-cold) for 1 minute to remove mucus as described (52). Planarians were fixed in 4% formaldehyde/1X PBS, or methacarn (6 ml methanol, 3 ml chloroform, 1 ml glacial acetic acid) for 10 min at room temperature as described (52). Formaldehyde-fixed samples were washed 3 times (5 minutes each) in 1X PBS. Methacarn-fixed samples were first rinsed 3 times in methanol, then rehydrated in 1:1 methanol:PBS for 5 minutes, followed by 3 washes (5 minutes each) in PBS. For analysis of RNA integrity, 5-10 planarians were immediately homogenized in Trizol, and RNA was extracted using two chloroform extractions and high-salt precipitation buffer according to the manufacturer’s instructions. RNA samples were analyzed using Agilent RNA ScreenTape on an Agilent TapeStation 2200 according to the manufacturer’s protocol.

For histology on methacarn-fixed samples, animals were relaxed and fixed individually in glass vials to minimize adherence to other samples. After fixation and rehydration, animals were incubated in 5%, 15%, and 30% sucrose (in RNAse-free 1X PBS), for 5-10 min each. Samples were then mounted and frozen in OCT medium, and cryosectioned at 20 μm thickness onto either Superfrost Plus glass slides (Fisher 12-550-15) (for staining optimization) or PEN membrane slides (Fisher/Leica No. 11505158) (for LCM). Prior to cryosectioning, PEN membrane slides were treated for 1 min with RNAse*ZAP* (Invitrogen AM9780), then rinsed by dipping 10 times (1-2 seconds per dip) in three successive conical tubes filled with 30 ml DEPC-treated water, followed by 10 dips in 95% ethanol and air-drying for 5-10 min. After cryosectioning, slides were stored on dry ice for 2-4 hours prior to staining. Cryostat stage and blades were wiped with 100% ethanol prior to sectioning.

For histological staining, slides were warmed to room temp. for 5-10 min. All slides were stained individually in RNAse-free conical tubes by manually dipping (1-2 seconds per dip) in 30 ml solutions. Forceps used for dipping were treated with RNAse*ZAP* and ethanol. All ethanol solutions were made with 200 proof ethanol and DEPC-treated water.

For Hematoxylin staining: 70% ethanol (20 dips); DEPC-treated water (20 dips); Mayer’s Hematoxylin (Sigma Aldrich MHS16-500ML) (15 dips); DEPC-treated water (10 dips); Scott’s Tap Water (2 g sodium bicarbonate plus 10 g anhydrous MgSO_4_ per liter of nuclease-free water) (10 dips); 70% ethanol (10 dips); 95% ethanol (10 dips); 95% ethanol (10 dips); 100% ethanol.

For Eosin Y staining: 70% ethanol (20 dips); DEPC-treated water (20 dips); 70% ethanol (20 dips); Alcoholic Eosin Y (Sigma Aldrich HT110116-500ML) (100%, 10%, or 2% diluted into 200 proof ethanol) (15 dips); 95% ethanol (10 dips); 95% ethanol (10 dips); 100% ethanol. In some cases fewer dips (8–10) in Eosin Y were required for better differentiation of gut tissue.

For combined Hematoxylin and Eosin Y staining: 70% ethanol (20 dips); DEPC-treated water (20 dips); Mayer’s Hematoxylin (15 dips); DEPC-treated water (10 dips); Scott’s Tap Water (10 dips); 70% ethanol (10 dips); 10% Alcoholic Eosin Y (15 dips); 95% ethanol (10 dips); 95% ethanol (10 dips); 100% ethanol.

The entire staining protocol was completed in less than 5 minutes. Slides were air dried for 5 minutes, then stored in plastic slide boxes on dry ice for 2-4 hr before LCM. Although we tested overnight storage at −80°C, we found that section morphology and RNA quality were best when conducting all steps, from fixation to LCM, on the same day.

### Laser-capture microdissection and RNA extraction

Stained PEN slides were removed from dry ice and immediately immersed for 30 seconds in ice-cold 100% ethanol, then room temperature 100% ethanol to minimize condensation/rehydration of sections and maintain RNAse inactivation during warming. Slides were then air dried for 2-3 minutes, and mounted in a Leica LMD7 laser microdissection microscope. Samples were dissected at 10X magnification using the following parameters: Power-30; Aperture-20; Speed-5; Specimen Balance-1; Head Current-100%; Pulse Frequency-392 Hz. Regions were dissected into empty RNAse-free 0.5 ml microcentrifuge caps (Axygen PCR-05-C). We separately captured medial intestine, lateral intestine, and non-intestine regions from all sections (8–10) per slide within 45-50 min. After capture, 20 μl Buffer XB (Arcturus PicoPure RNA Isolation Kit 12204-1) was added to captured tissue, then tubes were immediately frozen on dry ice and stored at −80°C prior to RNA extraction.

For RNA extraction, samples were thawed for 5 min at room temp. Next, tissue from two tubes/slides (16-20 sections from the same planarian) was pooled for each biological replicate, incubated at 42°C for 30 minutes, and RNA was extracted using the Arcturus PicoPure RNA Isolation Kit following the manufacturer’s instructions. 40-400 ng of total RNA was obtained from each sample, measured using a Denovix UV Spectrophotometer. RNA quality was analyzed using Agilent HS RNA ScreenTape on an Agilent TapeStation 2200 according to the manufacturer’s protocol.

### Library preparation and RNA sequencing

Concentration of RNA was ascertained using a Thermo Fisher Qubit fluorometer. RNA quality was verified using the Agilent Tapestation. Libraries were generated using the Lexogen Quantseq 3’ mRNA Library Prep Kit according to the manufacturer’s protocol, with 5 ng total RNA input for each sample. Briefly, first-strand cDNA was generated using 5’-tagged poly-T oligomer primers. Following RNase digestion, second strand cDNA was generated using 5’-tagged random primers. A subsequent PCR step with additional primers added the complete adapter sequence to the initial 5’ tags, added unique indices for demultiplexing of samples, and amplified the library. Final libraries for each sample were assayed on the Agilent Tapestation for appropriate size and quantity. Libraries were then pooled in equimolar amounts as ascertained by fluorometric analysis. Final pools were quantified using qPCR on a Roche LightCycler 480 instrument with Kapa Biosystems Illumina Library Quantification reagents. Sequencing was performed using custom primers on an Illumina Nextseq 500 instrument with High Output chemistry and 75 bp single-ended reads.

### Short-read mapping and gene-expression analysis

Adapters and low quality reads were trimmed from fastq sequence files with BBDuk (https://sourceforge.net/projects/bbmap/) using Lexogen data analysis recommendations (https://www.lexogen.com/quantseq-data-analysis/): k=13 ktrim=r forcetrimleft=11 useshortkmers=t mink=5 qtrim=t trimq=10 minlength=20. Sequence quality was assessed before and after trimming using FastQC (159). Reads were then mapped to a version of the *de novo* dd_Smed_v6 transcriptome (11) restricted to 28,069 unique transcripts (i.e., those transcripts whose identifiers ended with the suffix “_1”) using Bowtie2 (v2.3.1) (160) with default settings. Resulting SAM files were converted to BAM files, sorted, and indexed using Samtools (v1.3) (161). Raw read counts per transcript were then generated for each BAM file using the “idxtats” command in Samtools, and consolidated into a single Excel spreadsheet.

The resulting read counts matrix was imported into R, then analyzed in edgeR v3.8.6 (162). First, all transcripts with counts per million (CPM) < 1 in 4/12 samples (e.g., lowly expressed transcripts) were excluded from further analysis (13,136 / 28,069 transcripts were retained). Next, after recalculation of library size, samples were normalized using trimmed mean of M-values (TMM) method, followed by calculation of common, trended, and tagwise dispersions. Finally, differentially expressed transcripts in intestinal vs. non-intestinal samples were determined using the generalized linear model (GLM) likelihood ratio test. 1911 transcripts had a fold-change of more than 2 (logFC>1) and an FDR-adjusted *p* value <.01 in either medial or lateral intestine, relative to non-intestinal tissue. We further limited to 1844 transcripts with a minimum transcripts-per-million (TPM) of 2 in 4 of any 8 intestinal biological replicates (medial or lateral), since transcripts with lower expression values were at the lower limit of detection by ISH, and their removal also modestly increased the robustness of LCM vs. phagocyte analysis.

### Human Protein Atlas comparison

We queried (TBLASTX) 28,069 unique dd_Smed_v6 nucleotide sequences against 20,726 nucleotide sequences in the human reference proteome downloaded from UniProt (www.uniprot.org) (Release 2017_12, 20-Dec-2017). 13,362 dd_Smed_v6 transcripts hit human sequences (7,309 unique) with *E*-value ≤ 1 x 10^-3^. These human sequences were then used to conduct reciprocal TBLASTX queries against dd_Smed_v6 transcripts: 7,220 hit 5,808 unique dd_Smed_v6 sequences with *E*-value ≤ 1 x 10^-3^. In total, 5,583 dd_Smed_v6 transcripts had reciprocal best hits (RBHs) in the human proteome, with >94% having *E*-value ≤ 1 x 10^-10^ in either direction. Of 1844 intestine-enriched transcripts, 701 had RBHs in the human proteome.

Next, using RBH UniProt Accession numbers, we extracted RNA-Seq tissue enrichment data from the Human Protein Atlas (95). 699/701 intestinal transcripts’ RBH homologs were present in the HPA data (Table S4); of these, 130 were enriched in one or more human tissues. 5,561/5,583 dd_Smed_v6 transcripts’ RBH homologs were present in the HPA data; of these, 1,011 were enriched in one or more human tissues. The number of intestine and non-intestine RBH homologs enriched in each of 32 human tissues was calculated, and expressed as a percentage of total transcripts (699 intestine or 5,561 non-intestine) with RBH homologs. Finally, ratio of the percentage of intestine-enriched RBH homologs to the percentage of all dd_Smed_v6 RBH homologs was then calculated to determine fold-enrichment for each tissue. Two tissues (“Appendix” and “Smooth Muscle”) were excluded from final analysis since <0.1% (6/5561) of all dd_Smed_v6 transcripts had RBH homologs enriched in these tissues.

TBLASTX queries were conducted using NCBI BLAST+ standalone suite. Extraction and analysis of tissue enrichment data from HPA was conducted in R and Excel.

### Gene Ontology annotation, nomenclature, and analysis

BLASTX homology searches of UniProtKB protein sequences for *H. sapiens, M. musculus, D. rerio, D. melanogaster,* and *C. elegans* were conducted using all 28,069 unique transcripts in the “dd_Smed_v6” transcriptome in PlanMine as queries. In all figures, gene names/abbreviations are based on the best (lowest *E*-value) UniProt homolog, except: (1) genes were named after the best human homolog for HPA analysis (Figure 3); and (2) we used *Smed* nomenclature when genes (or paralogs) were previously named by us or others, including *hnf-4/*dd_1694_0_1 (33), *nkx2.2*/dd_2716_0_1 (24), *gata4/5/6/*dd_4075_0_1 (33), *apob-1/*dd_636_0_1 [DJF, unpublished], *apob-2/*dd_194_0_1 [DJF, unpublished], *slc22a6*/dd_1159_0_1 (65), *gli-1/*dd_7470_0_1 (108), and *RREB2/*dd_10103_0_1 (141). Biological Process GO terms (also obtained from UniProtKB) from the top hit for each species (with *E*-value ≤ 1 x 10^-5^) were assigned to each dd_Smed_v6 transcript. 9344 of 28,069 total transcripts and 1379 of 1844 intestine-enriched transcripts were annotated with GO terms. Enrichment for specific terms among all intestine-enriched transcripts, medially enriched transcripts, or laterally enriched transcripts was then evaluated with BiNGO (163). Regional transcript enrichment was calculated as a ratio of fold changes in medial and lateral intestinal tissue: transcripts were considered to be medially enriched if FC_medial_/FC_lateral_>1 (1221 transcripts), or laterally enriched if FC_medial_/FC_lateral_<1 (623 transcripts). All 13,136 (of 28,069) transcripts detected in our experiment (above) were used as a background set, and hypergeometric testing with a Benjamini & Hochberg False Discovery Rate (FDR) of .05 was considered significant. We additionally restricted our summarization to terms that were annotated to more than one percent of transcripts that received annotations in each group: 1379/1844 intestine-enriched transcripts, 924/1221 medially enriched transcripts, or 455/623 laterally enriched transcripts.

### Comparison to gene sets involved in innate immunity

1456 *Dugesia japonica* transcripts upregulated in response to either *L. pneumophila* or *S. aureus* infection (91) were queried (TBLASTX) against 28,069 dd_Smed_v6_unique transcripts. 981 dd_smed_v6 transcripts hit with evalues < 1e-03. After removing duplicate hits, 783 transcripts remained. 701/783 transcripts were present in the full 13,136 LCM dataset; 99/783 were intestine-enriched.

30,021 SMED_20140614 transcripts were mapped to 28,069 dd_smed_v6_unique transcripts, generating 22,889 dd_smed_v6_unique hits with evalues <1e-03. After removing duplicates, 19,338 transcripts remained. 741/19,338 transcripts were up- or down-regulated in planarians shifted to static culture (90). 447/741 transcripts were present in the full 13,136 LCM dataset, and 62 were intestine-enriched.

### Comparison to gene expression in sorted phagocytes

28,069 unique dd_Smed_v6 transcripts were blasted (BLASTN) against 11,589 ESTs and assembled contigs from the “SmedESTs3” collection (164), which were used in microarray-based analysis of gene expression in sorted phagocytes (24). 7927 hit with length >100 bp, >90% base identity, and *E-*value less than 1 x 10^-50^. Of these, 6919 of 13,136 Smed_v6 transcripts detected in LCM samples mapped to unique Smed_ESTs3 contigs or ESTs with detectable expression in the sorted phagocyte data set. 2626/6919 transcripts had FDR-adjusted *p* values <.01 for logFC in either medial or lateral intestine (vs. non-intestine, LCM data in this study), and 1498/2626 had FDR-adjusted *p* values <.05 for logFC in sorted phagocytes (vs. all other cells) (phagocyte data in (24)). 1317/2626 had logFC >1 in either medial or lateral intestine. In the sorted phagocyte data, 900/1317 had a fold-change >0 and FDR-adjusted *p* value <.05 (“high” in phagocytes), 358/1317 had an FDR-adjusted *p* value >.05 (“moderate” in phagocytes), and 59/1317 had a fold-change <0 and FDR-adjusted *p* value >.05 (“low” in phagocytes).

### Comparison to gene expression in single-cell transcriptomes

We utilized data from gene expression analysis of single planarian cells from two recent studies (32, 43) to identify dd_Smed_v6 transcripts enriched in specific intestinal cell types that were also represented in our 1844 intestine-enriched transcripts (this study) and phagocyte expression data (24). For comparison to Fincher et al. (32) (Fig S4D-F), we identified 391 phagocyte-enriched transcripts (subcluster 4 or “enterocytes” in Table S2 (intestine) (32)), 21 goblet-enriched transcripts (subcluster 8 in (32)), and 30 basal-enriched transcripts (subcluster 8 or “outer intestinal cells” in (32)); transcripts found in other intestinal subclusters were excluded. For comparison to Plass et al. (43) (Fig S4G-H), we identified 92 phagocyte-enriched transcripts and 27 goblet cell-enriched transcripts that were also represented in our 1844 intestine-enriched transcripts and phagocyte expression data. For global comparison of transcripts enriched in laser captured intestine (Fig S4I), we included all transcripts enriched in intestinal cell types (and intestinal precursors) in single cell studies (32, 43, 88), without regard to cell type or enrichment in non-intestinal cell types. For comparison to Swapna et al. (88) data, BLASTN was used to identify dd_Smed_v6 transcripts orthologous to “SmedASXL” transcripts (with >95% identity and length >100 bp).

### Gene cloning and Expressed Sequence Tags

Total RNA was extracted from planarians using Trizol with two chloroform extractions and high salt precipitation. After DNAse digestion, cDNA was synthesized using the iScript Kit (Bio-Rad 1708891). Genes were amplified by PCR with Platinum *Taq* (Invitrogen 10966026) using primers listed in Table S1C. Amplicons were cloned into pJC53.2 digested with Eam1105I as described (130) and sequenced to verify clone identity and orientation. For some genes, expressed sequence tags (ESTs) in pBluescript II SK(+) were utilized (164), also listed in Table S1C.

### In situ hybridization and immunofluorescence

In situ hybridizations were performed as described (60), with the following adjustments: NAc (7.5%) treatment was for 15 min; 4% formaldehyde fixation was for 15 min; and animals were bleached for 3 hr. Samples were pre-incubated with tyramide solution without H_2_O_2_ for 10 min, then spiked with H_2_O_2_ (.0003% final concentration), then developed for 10 min. Immmunofluorescence after FISH was conducted as described (53), using 1x PBS, 0.3% Trition-X100, 0.6% IgG-free BSA, 0.45% fish gelatin as the blocking buffer (52).

### Mucin identification

*S. mediterranea* mucin-like genes in PlanMine (11) and SmedGD2.0 (10) were identified by TBLASTN searches with human refseq_protein and UniProt mucin sequences. Planarian sequences were translated with NCBI ORFinder (https://www.ncbi.nlm.nih.gov/orffinder/), and domain searches were conducted using NCBI CD-Search (https://www.ncbi.nlm.nih.gov/Structure/cdd/cdd.shtml) (165), Pfam 31.0 (https://pfam.xfam.org) (166), and SMART (http://smart.embl-heidelberg.de) (167). Three planarian sequences were identified that encoded three N-terminal von Willebrand factor D domains and two or more N-terminal cysteine-rich and trypsin-inhibitor-like cysteine-rich domains characteristic of human mucins (e.g., MUC-2, MUC-5AC) (168, 169), but not von Willebrand factor A or Thrombospondin type I repeats found in closely related proteins (e.g., SCO-spondin). Planarian mucin-like genes identified using this approach are: *Smed-muc-like-1* (dd_Smed_v6_17988_0_1/SMED30009111), *Smed-muc-like-2* (dd_Smed_v6_18786_0_1/SMED30002668), and *Smed-muc-like-3* (dd_Smed_v6_21309_0_1/dd_Smed_v6_38233_0_1/dd_Smed_35076_0_1/SMED30006765).

### Identification of intestine-enriched transcription factors and RNA interference

Putative transcription factors were identified by extracting intestine-enriched transcripts with “DNA” and/or “transcription” Biological Process and Molecular Function GO terms, then verifying that the best Uniprot homologs regulated transcription in published experimental evidence. *Smed-gli-1* has been previously studied (108). *Smed-RREB2* was most homologous to another planarian zinc finger protein, *Smed-RREBP1* (141). However, we used the more common “*RREB*” abbreviation (rather than “*RREBP*”) for the second planarian paralog.

RNAi experiments were conducted as described (170) by mixing one microgram of in vitro-synthesized dsRNA with 1 μl of food coloring, 8 μl of water, and 40 μl of 2:1 liver:water homogenate. For the primary regeneration screen, animals were fed three times over 6 days, amputated 4-5 days after the last feeding, then fixed 6 days post amputation. For further analysis of *gli-1(RNAi)* and *RREB2(RNAi)* phenotypes in uninjured animals, planarians were fed 1X/week for 6 weeks or 2X/week for 3 weeks (6 feedings total). *gli-1(RNAi)* and *RREB2(RNAi)* animals were scored as “non-eaters” if they refused food on two successive days (7 and 8 days after the previous feeding). Non-eating and/or curling animals were excluded from FISH analysis. For experiments in regenerates, animals were fed 2X/week for 4 weeks (8 feedings total).

### Image Collection

Confocal images were collected on a Zeiss 710 Confocal microscope, with the following settings: two tracks were used, one with both 405/445 (excitation/emission nm, blue) and 565/650 (red), and the other with only 501/540 (green) to minimize bleed-through between channels. Detector gains were adjusted so that no pixels were saturated. Digital offset was set to 0 or −1, to ensure that most pixel intensities were non-zero, with averaging set to 2. Whole animals were captured with a single z-plane (5 µm section) with a 10x objective, tiled, and stitched in Zen (version 11.0.3.190, 2012-SP2). Magnified regions were captured with a 63x Oil immersion objective and a z-stack (50-100 slices, 0.31 µm/slice) was collected from the tail and/or head of the animal. In some cases, after imaging min/max or linear best fit adjustments were made in Zen to improve contrast of final images. Profile graphs (Figs 5-7) represent raw, unadjusted pixel intensity values from a single optical section.

WISH images were collected on a Zeiss Stemi 508 with an Axiocam 105 color camera, an Olympus SZX12 dissection microscope with an Axiocam MRc color camera, or a Zeiss Axio Zoom.V16 with an Axiocam 105 camera. In some cases brightness and/or contrast were adjusted in photoshop to improve signal contrast.

### Quantification of animal area and length for *gli-1* and *RREB2* knockdown animals

Animals were separated into 35mm x 10mm petri dishes in groups of two, and imaged on a Zeiss Stemi 508 prior to the first dsRNA feeding, then again seven days after the fifth feeding (animals were fed once per week). For area, images were processed with ImageJ by first applying an Auto Threshold, using method = Intermodes, Huang, or Triangles with “White objects on black background” selected, to highlight the planarian. Analyze Particles was then run, with size = 600000-10000000 µm^2^, and “Display results” and “Include holes” selected. For length, a straight line was drawn manually from the tip of the head to the tip of the tail and then Measure was used to obtain the length.

### Data availability

Raw and processed RNA-Seq data associated with this study have been deposited in the NCBI Gene Expression Omnibus (GEO) under accession number GSE135351.

## Acknowledgments

We thank Forrest Waters for initial optimization of fixation, labeling, and RNA extraction for laser microdissection. We thank Mayandi Sivagaru (Institute for Genomic Biology, University of Illinois at Urbana-Champaign) and Muralidharan Jayaraman (University of Oklahoma Health Sciences Center) for help with LCM optimization, and Alvaro Hernandez (Roy J. Carver Biotechnology Center, Illinois) and Graham Wiley (OMRF Clinical Genomics Center) for library preparation and Illumina sequencing. We thank Ben Fowler, Amanda Valdez, Julie Crane, and Alex Weddle (OMRF Imaging Core) for technical assistance with confocal microscopy, histology, and LCM slide preparation; Jonathan Wren (OMRF) (Bioinformatics and Pathways Core) for advice on reciprocal blast searches; and Stuart Glenn and the OMRF Quantitative Analysis Core for high performance computing resources. We are grateful to Ricardo Zayas and Francesc Cebrià, who originally identified *Smed-npc2* as a marker for goblet cells. We gratefully acknowledge members of the Newmark and Forsthoefel Labs, Linda Thompson, Dean Dawson, and David Jones for discussions and manuscript critique.

## Supplementary Material Legends

**Figure S1.**
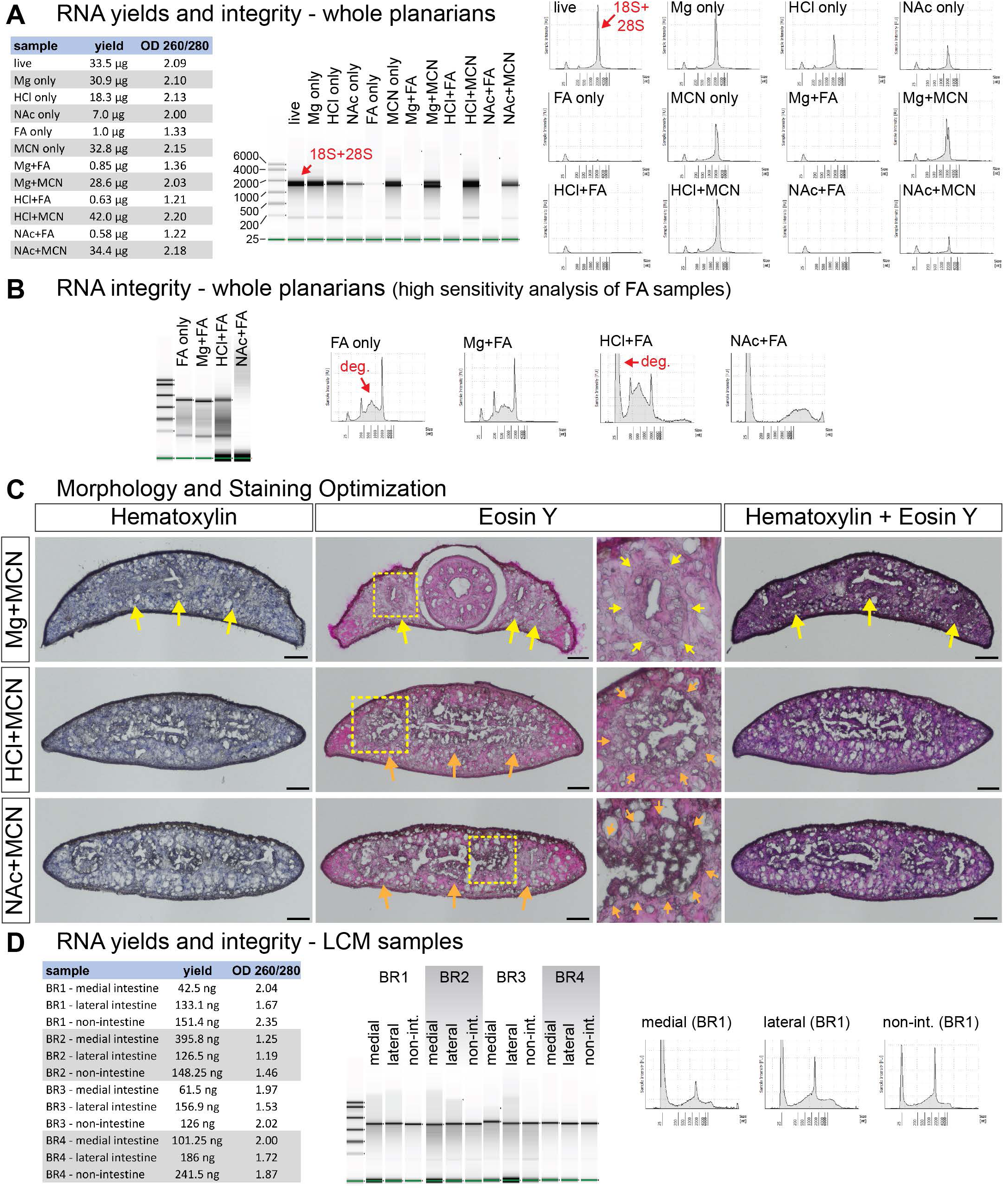
Optimization of fixation and histological staining for laser microdissection. **(A)** RNA yields (left table) and Bioanalyzer/TapeStation analysis of RNA integrity (right panels) from whole planarian samples (15 planarians, ∼20-25 mg total tissue) treated as shown. As in some other invertebrates (54–57), heat denaturation prior to analysis causes 28S rRNA to co-migrate as two bands with 18S rRNA (indicated with red arrows). Samples are representative of at least three independent replicates. RNA from FA-fixed samples had low yields and low 260/280 ratios. **(B)** Analysis of degraded RNA from FA-fixed samples using high-sensitivity (HS) TapeStation kit (for low concentration samples). Degradation of rRNA is indicated with red arrows. **(C)** Hematoxylin and Eosin Y labeling of transverse (cross) cryosections from methacarn-fixed planarians. Yellow arrows indicate intestine labeling in magnesium-treated samples. Orange arrows indicate compromised intestinal morphology in HCl- and NAc-treated samples. Boxed regions for Eosin Y-labeled sections are magnified to the right. **(D)** RNA yields (left table) and HS TapeStation analysis of RNA integrity (right panels) for all LCM samples used in this study (BR, biological replicates, 16-20 tissue sections per replicate). TapeStation lanes (middle panel) were assembled from multiple runs. Intensity plots (right panel) for BR1 are representative of other replicates. Mg, MgCl_2_ treatment. HCl, 2% HCl treatment. NAc, 7.5% N-Acetyl-L-Cysteine treatment. FA, 4% formaldehyde fixation. MCN, methacarn fixation. Scale bars: 100 μm (C).

**Figure S2.**
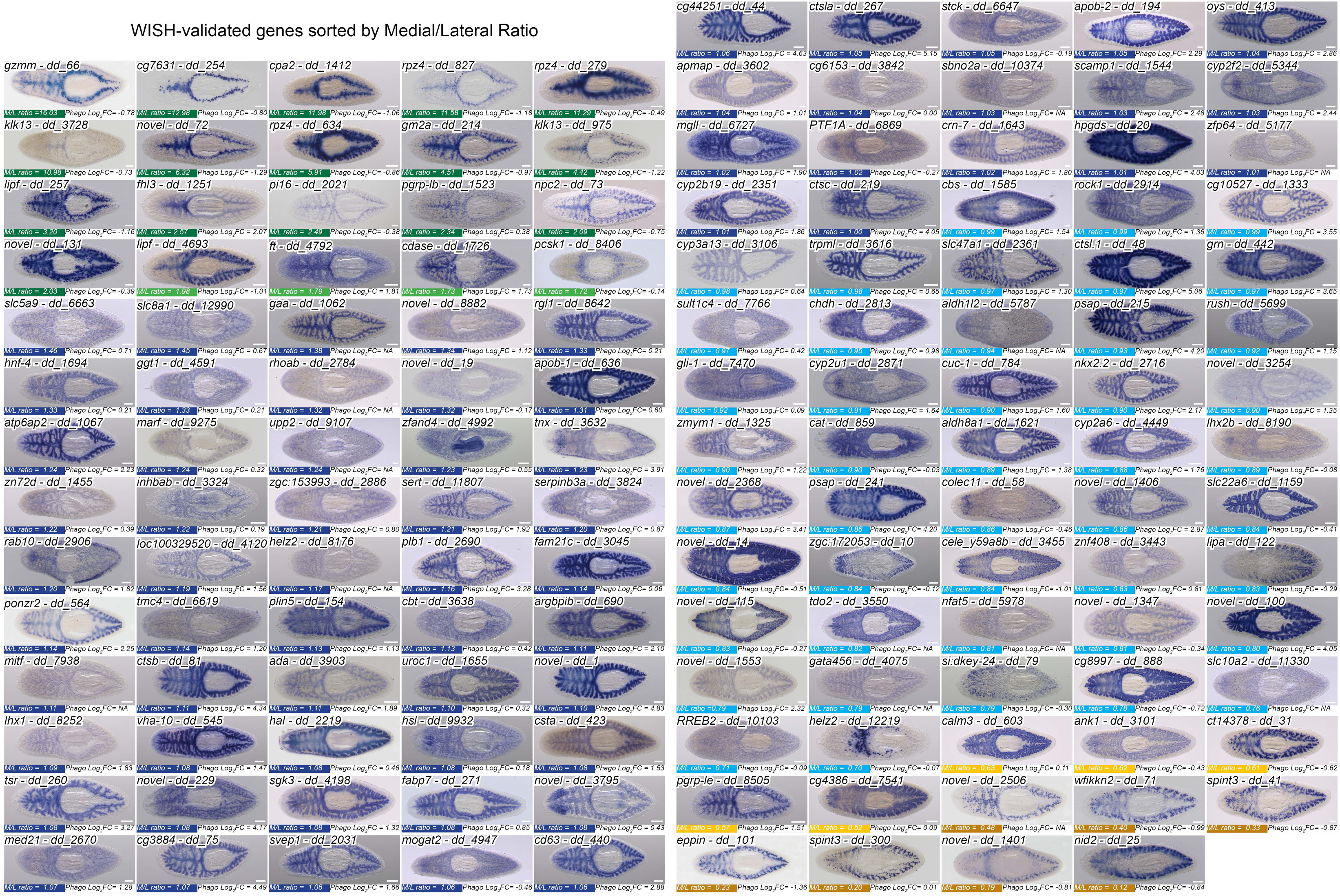
Expression of transcripts enriched in laser captured intestinal tissue. Whole-mount in situ hybridizations performed during this study, organized by mediolateral ratio. 144/162 total in situs conducted are shown (18 omissions were due to very low or no detectable expression; *helz2/*dd_12219 was detected only in peripharyngeal cells). Each image also shows the phagocyte enrichment value (Log_2_FC in Table S5), as well as the Uniprot best hit ID (Table S1) and the Dresden v6 Transcriptome GeneID (dd_Smed_v6). Several transcripts are named to be consistent with previous publications (*hnf-4/dd_1694, PTF1A/dd_6869, gli-1/dd_7470, slc22a6/dd_1159*), with human Uniprot IDs used in our comparison to the Human Protein Atlas (*sult1c4/dd_7766, tdo2/dd_3550*), or to distinguish paralogs (*apob-1/dd_636, apob-2/dd_194).* Detailed gene ID information is available in Table S1. Scale bars, 200 μm.

**Figure S3.**
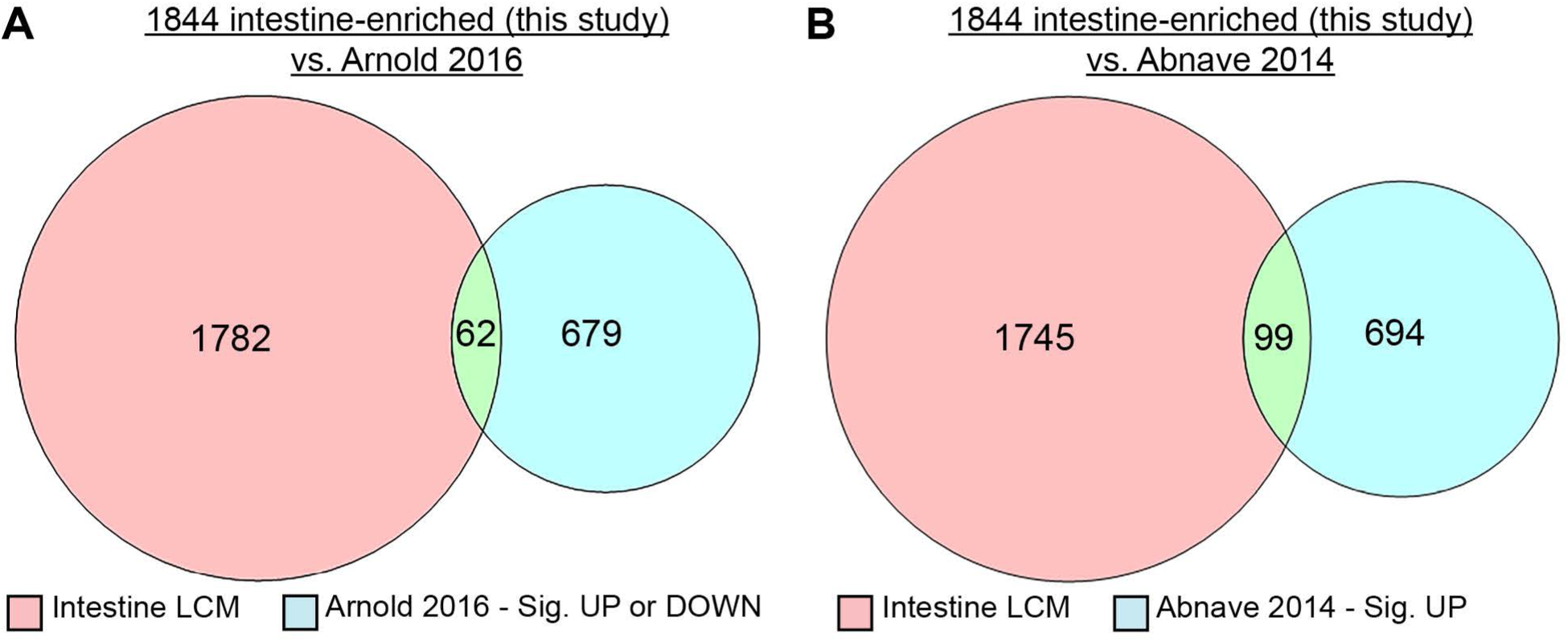
Innate immunity-related transcripts enriched in the intestine. **(A)** Intestine-enriched transcripts detected by LCM (pink) compared to *S. mediterranea* transcripts that were up- or downregulated upon transfer of planarians from continuous flow to static culture in Arnold et al., 2016 (90) (blue). **(B)** *S. mediterranea* intestine-enriched transcripts detected by LCM (pink) compared to *D. japonica* homologous transcripts that were upregulated in response to *L. pneumophila* or *S. aureus* infections in Abnave et al., 2014 (91) (blue).

**Figure S4.**
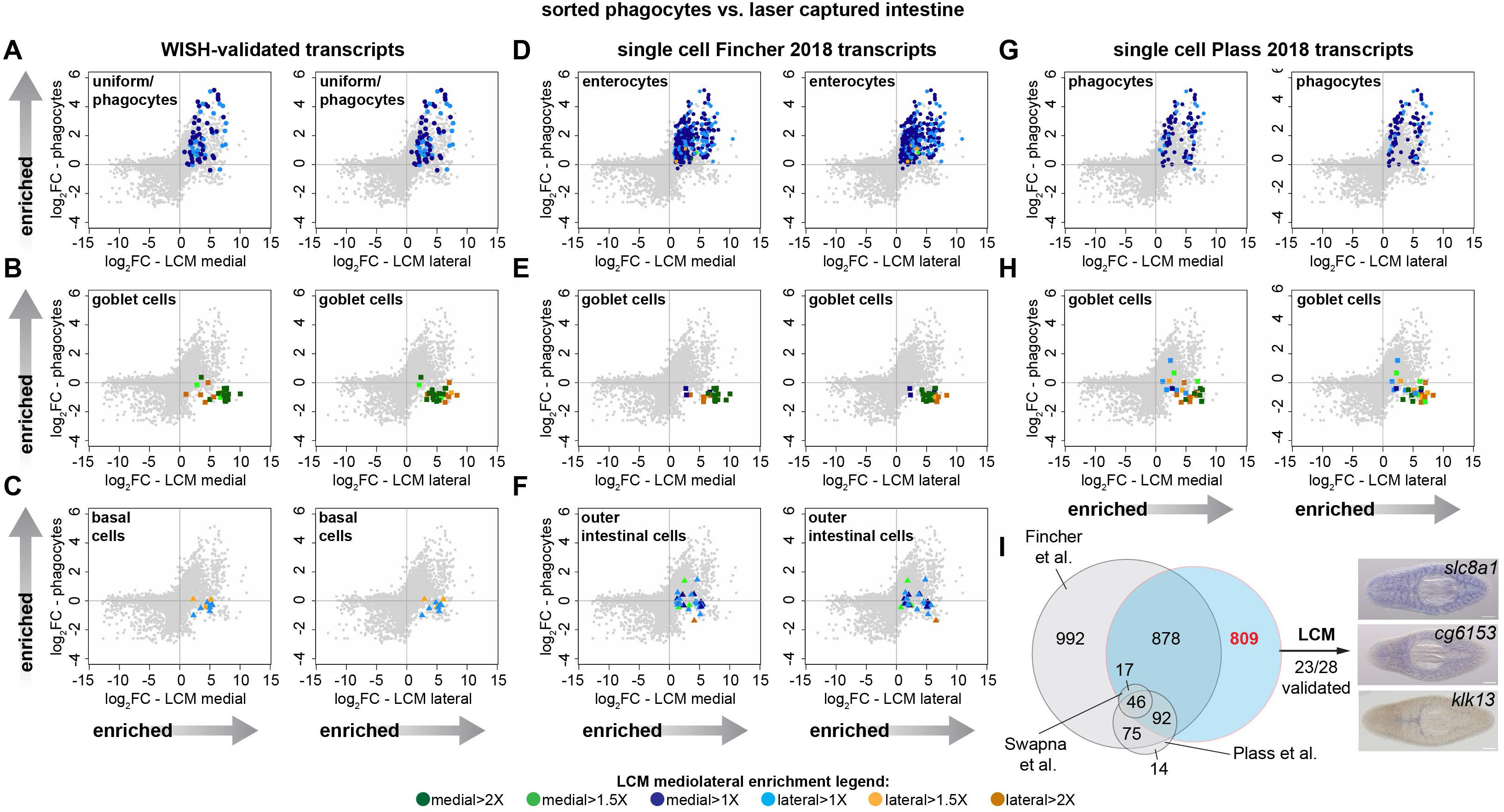
Validation of cell-type specific mRNA expression by WISH and correlation with single cell analysis. **(A-C)** Transcripts with WISH patterns indicating enrichment in phagocytes (A), goblet-like cells (B), or basal cells (C), mapped onto plots of phagocyte (y-axis) vs. LCM medial/lateral (x-axis) fold-change expression (log_2_FC). **(D-F)** Transcripts found to be highly enriched in phagocytes (“enterocytes”) (D), goblet cells (E), or basal cells (“outer intestinal cells”) in Fincher et al., 2018 (32), mapped onto phagocyte vs. LCM intestine expression plots. **(G-H)** Transcripts found to be highly enriched in phagocytes (G) or goblet cells (H) in Plass et al., 2018 (43), mapped onto phagocyte vs. LCM intestine expression plots (basal cells were not described). **(I)** Venn Diagram showing overlap between intestine-enriched transcripts in three recent single cell studies (32, 43, 88) and our LCM data. Overlaps of two or fewer genes are not displayed. Examples of 23/28 transcripts detected by LCM whose intestinal expression was validated by WISH. Detailed gene ID information is available in Table S1 and the Results. Scale bars, 200 μm.

**Figure S5.**
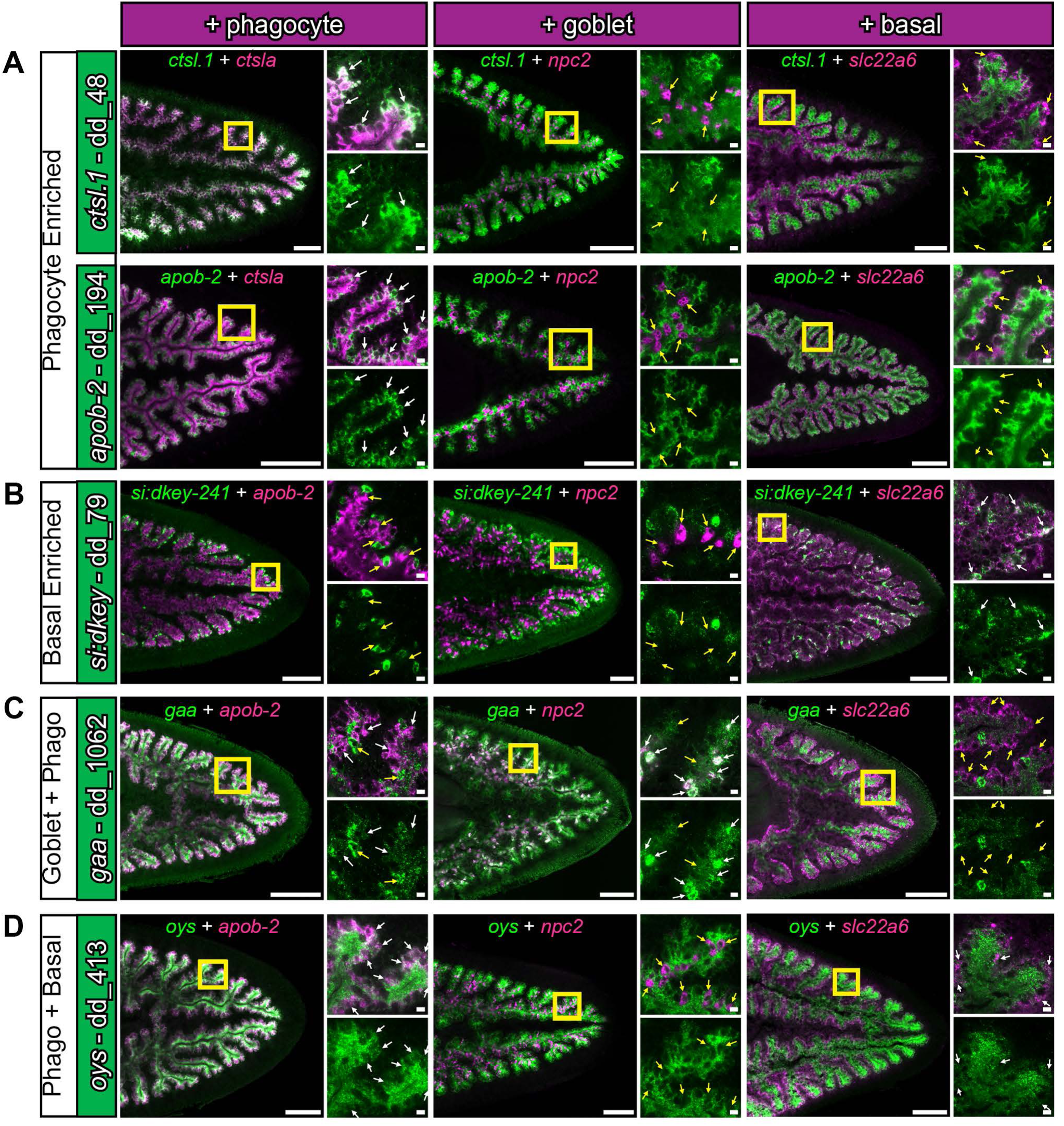
Additional cell-type specific transcripts revealed by double fluorescent in situ hybridization. **(A)** Phagocyte-enriched genes *ctsl.1* and *apob-2* (green), in combination with a phagocyte, goblet, or basal marker shown in Fig 5 (magenta). Yellow box indicates magnified region. White arrows indicate regions of considerable overlapping expression, while yellow arrows indicate regions of minimal overlap. **(B)** *si:dkey-241* (green) is enriched in a subset of *slc22a6-*positive basal cells. **(C)** *gaa* (green) is expressed in both goblet cells (high) and phagocytes (moderate), but is largely absent in basal cells. **(D)** *oys* is expressed in both phagocytes and basal cells. Detailed gene ID information is available in Table S1. Scale bars, whole animals 200 μm; magnified images, 10 µm.

**Figure S6.**
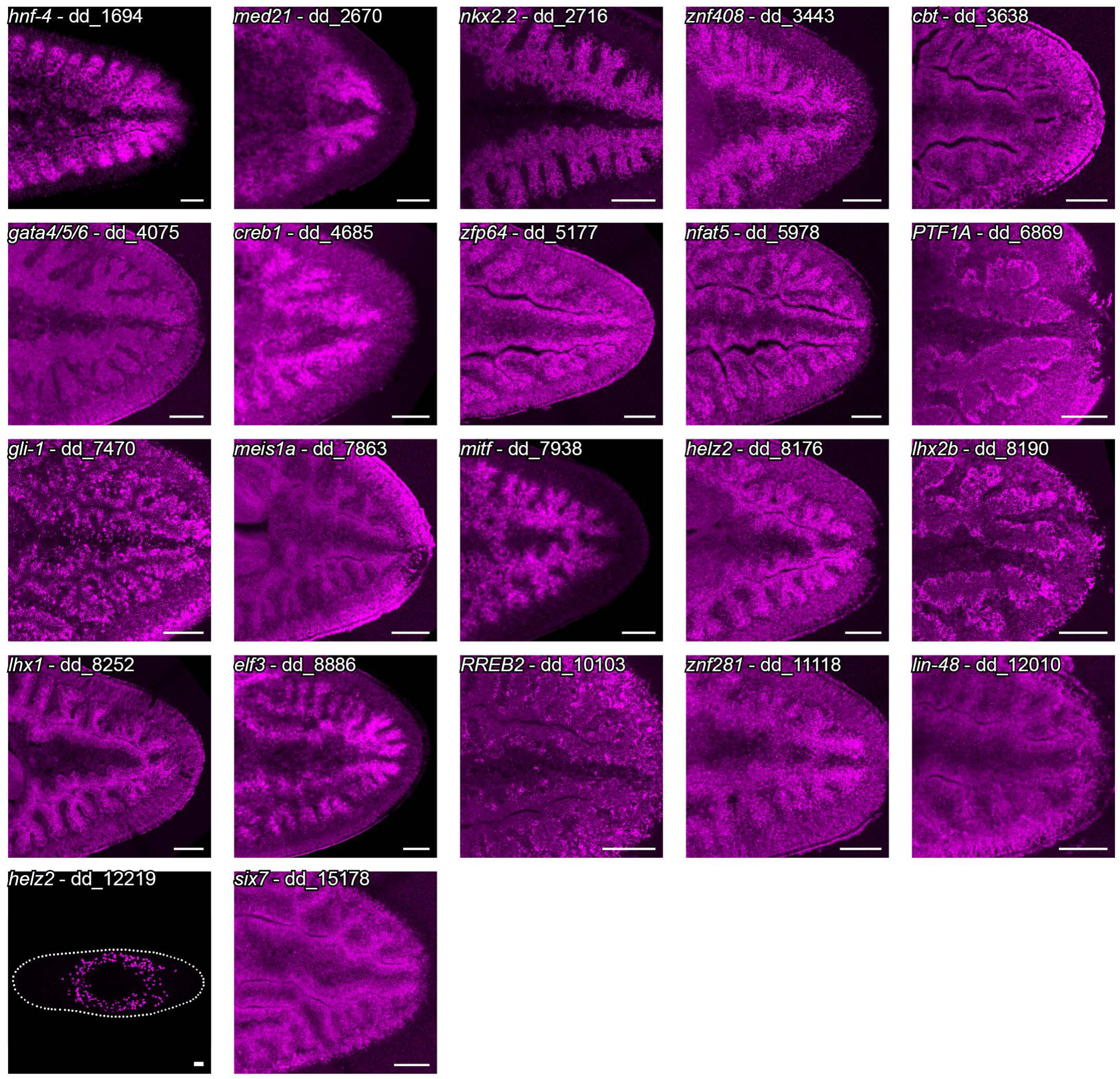
Expression of intestine-enriched transcription factors. Fluorescent in situ hybridization expression patterns of 22 intestine-enriched transcription factors identified in the laser capture dataset. Organized by dd_Smed_v6 ID in ascending order. Detailed gene ID information is available in Table S1. Scale bars, 100 μm.

**Figure S7.**
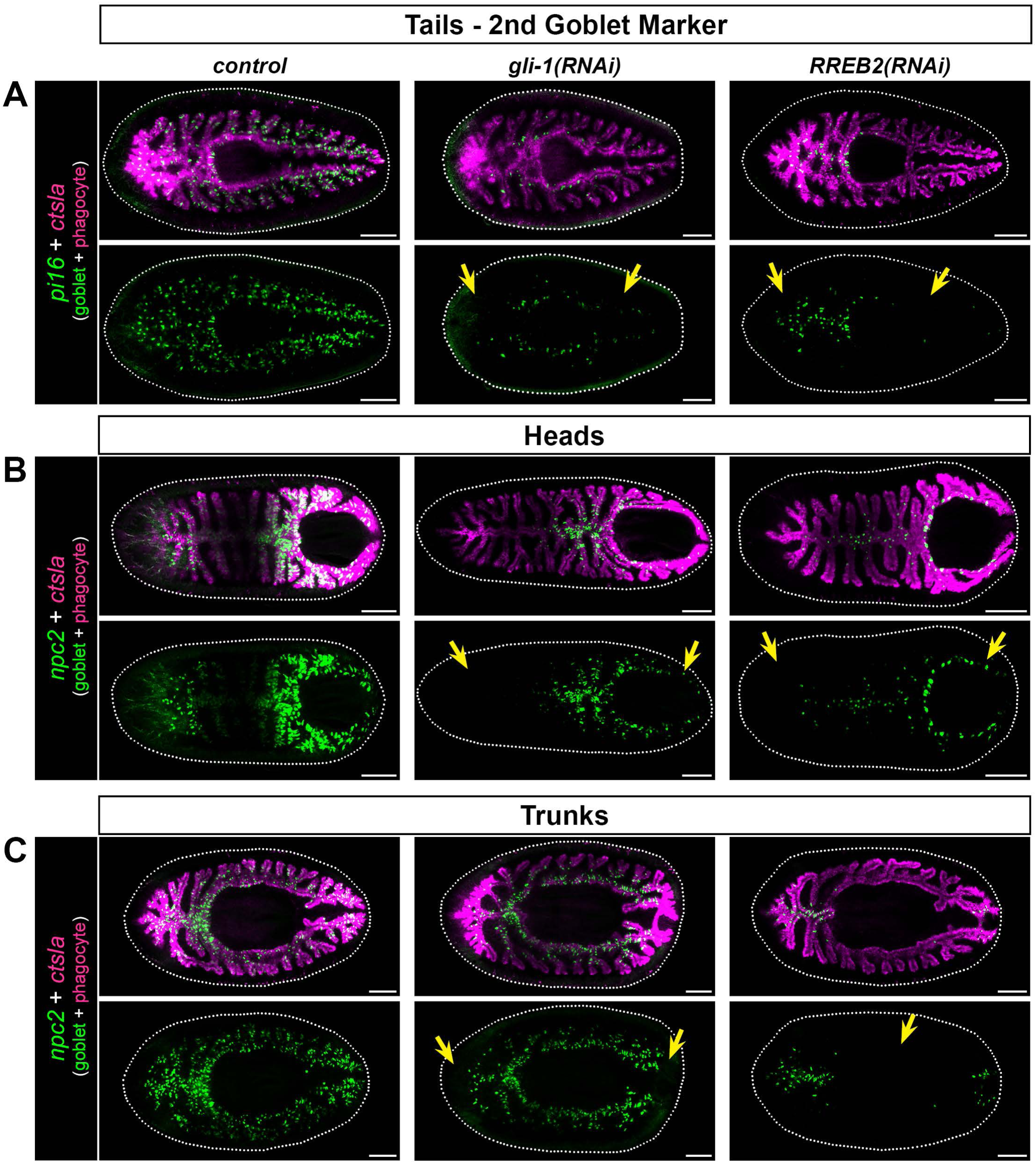
Goblet cells are reduced in *gli-1* and *RREB2* 7-day regenerates. Yellow arrows indicate regions of reduced or missing goblet cells. **(A)** Expression of a second, additional pan-goblet cell marker, *pi16*, in 7-day tail regenerates, which also shows dramatic reduction in goblet cell numbers (arrows). **(B)** *npc2* expression in 7-day regenerate heads, indicating severe reduction of goblet cells in both *gli-1 and RREB2* knockdowns (arrows). **(C)** Trunk fragments also show goblet cell loss. *gli-1* RNAi causes missing or reduced *npc2* expression in blastema regions, while *RREB2* reduces *npc2* labeling in pre-existing intestine (arrows). Detailed gene ID information is available in Table S1 and in Results. Scale bars, 200 μm.

**Figure S8.**
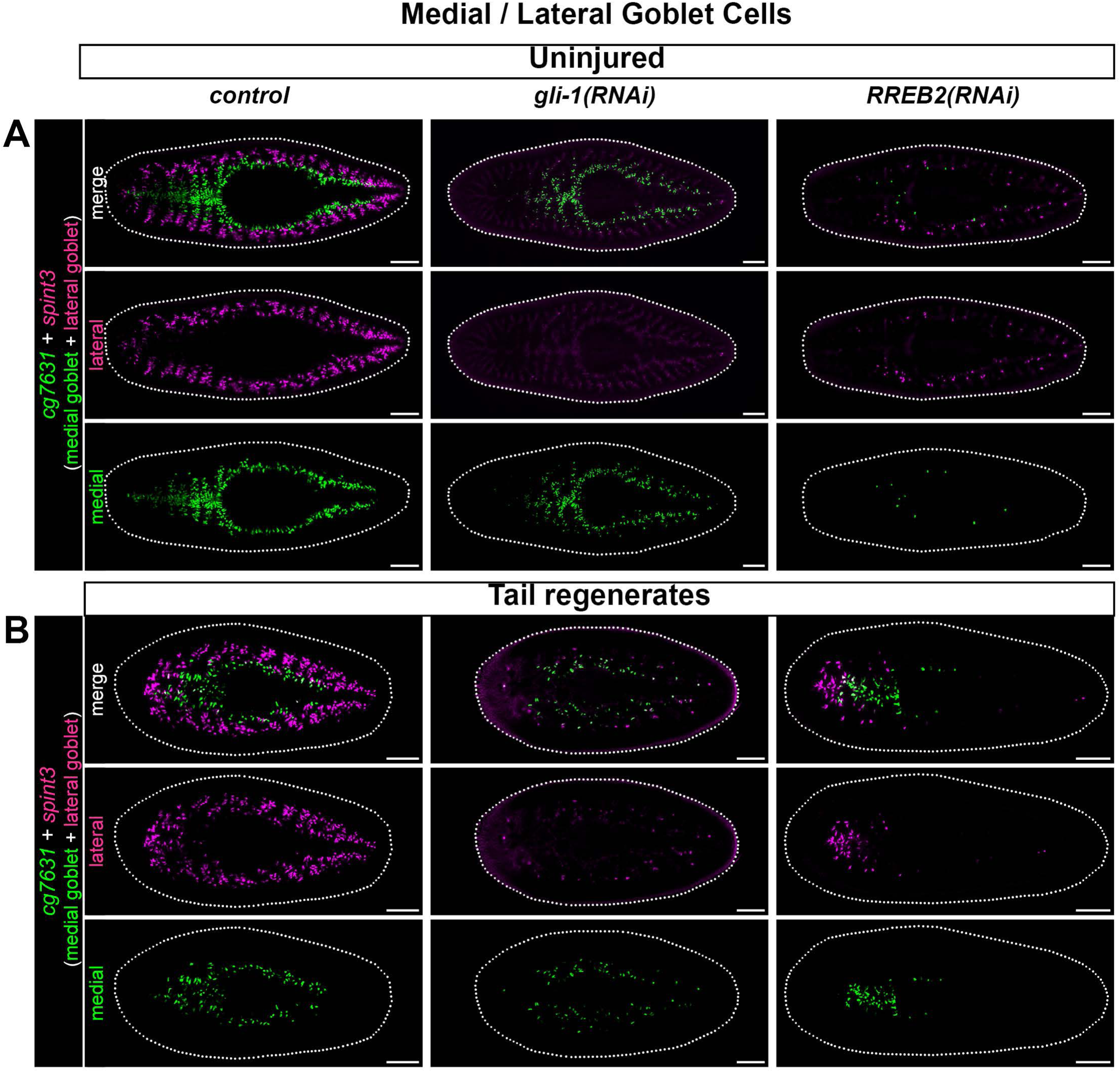
Effects of *gli-1* and *RREB2* knockdown on medial and lateral goblet cell subpopulations. **(A)** Uninjured animals showing the expression of *cg7631* (medial goblet) and *spint3* (lateral goblet) in control, *gli-1*, or *RREB2* knockdown animals. *gli-1(RNAi)* animals preferentially lose lateral goblet cells. *RREB2(RNAi)* animals lose both goblet cell subtypes, although more lateral cells remain compared to *gli-1* knockdown. **(B)** 7-day tail regenerates with medial and lateral goblet cells labeled in control, *gli-1*, or *RREB2* knockdowns. *gli-1* RNAi again reduces lateral goblet cells throughout the fragment, as well as medial populations in the anterior regenerating intestine, while *RREB2(RNAi)* regenerates lack all goblet cells in pre-injury intestine (e.g., posterior in tail fragments). Detailed gene ID information is available in Table S1. Scale bars, 200 μm.

**Figure S9.**
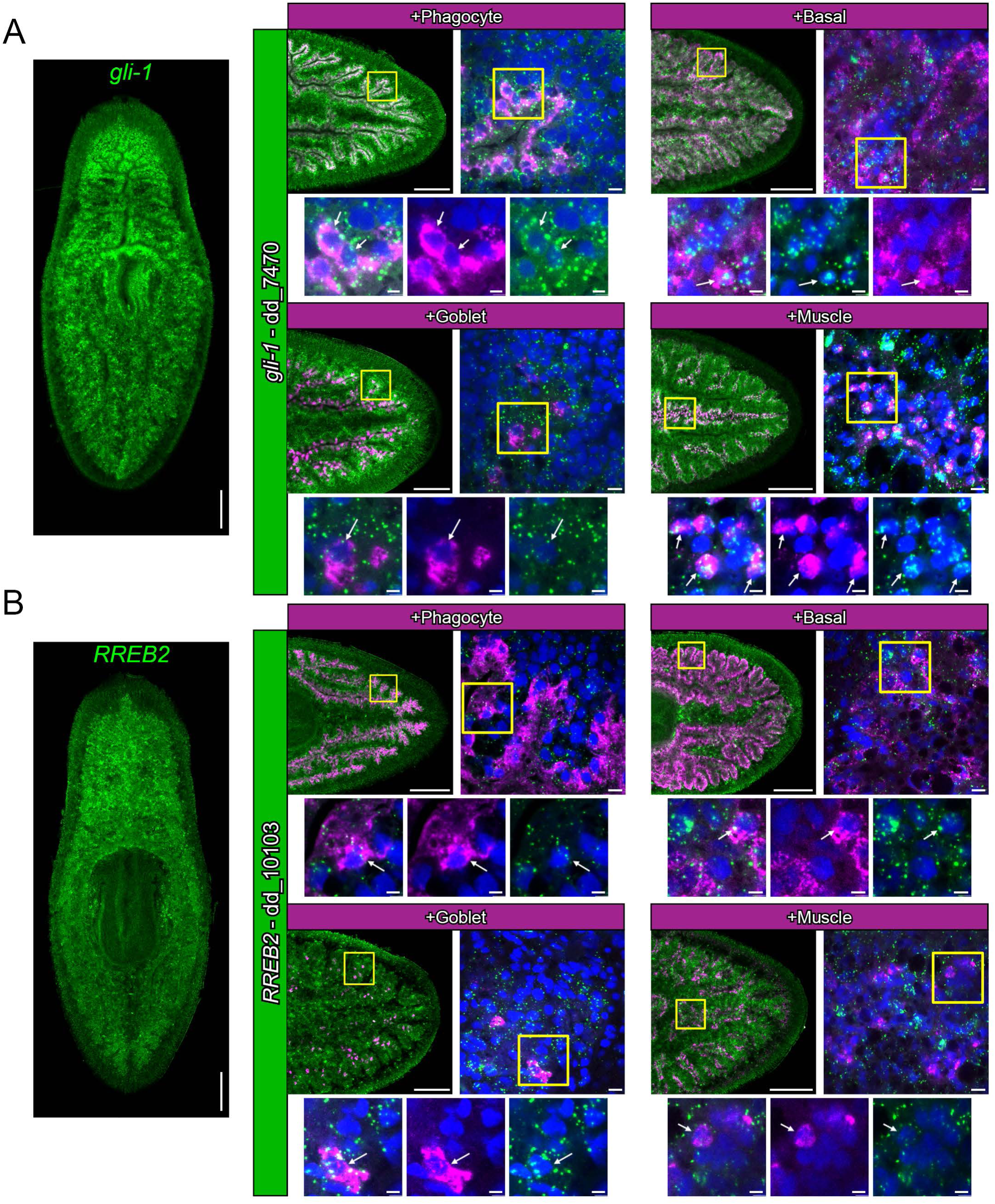
Expression patterns of *gli-1* and *RREB2*. **(A)** Global expression of *gli-1* in uninjured animals, along with images of mRNA expression in phagocytes (*ctsla*), basal cells (*slc22a6*), goblet cells (*npc2*), and muscle cells (*tni-4*). **(B)** Expression of *RREB2* in uninjured animals, along with mRNA co-localization with the same phagocyte, basal, goblet, and muscle cell markers as in A. Detailed gene ID information is available in Table S1. Scale bars, whole animals and tails 200 µm magnified images, 10 µm; digital zooms, 5 µm.

**Figure S10.**
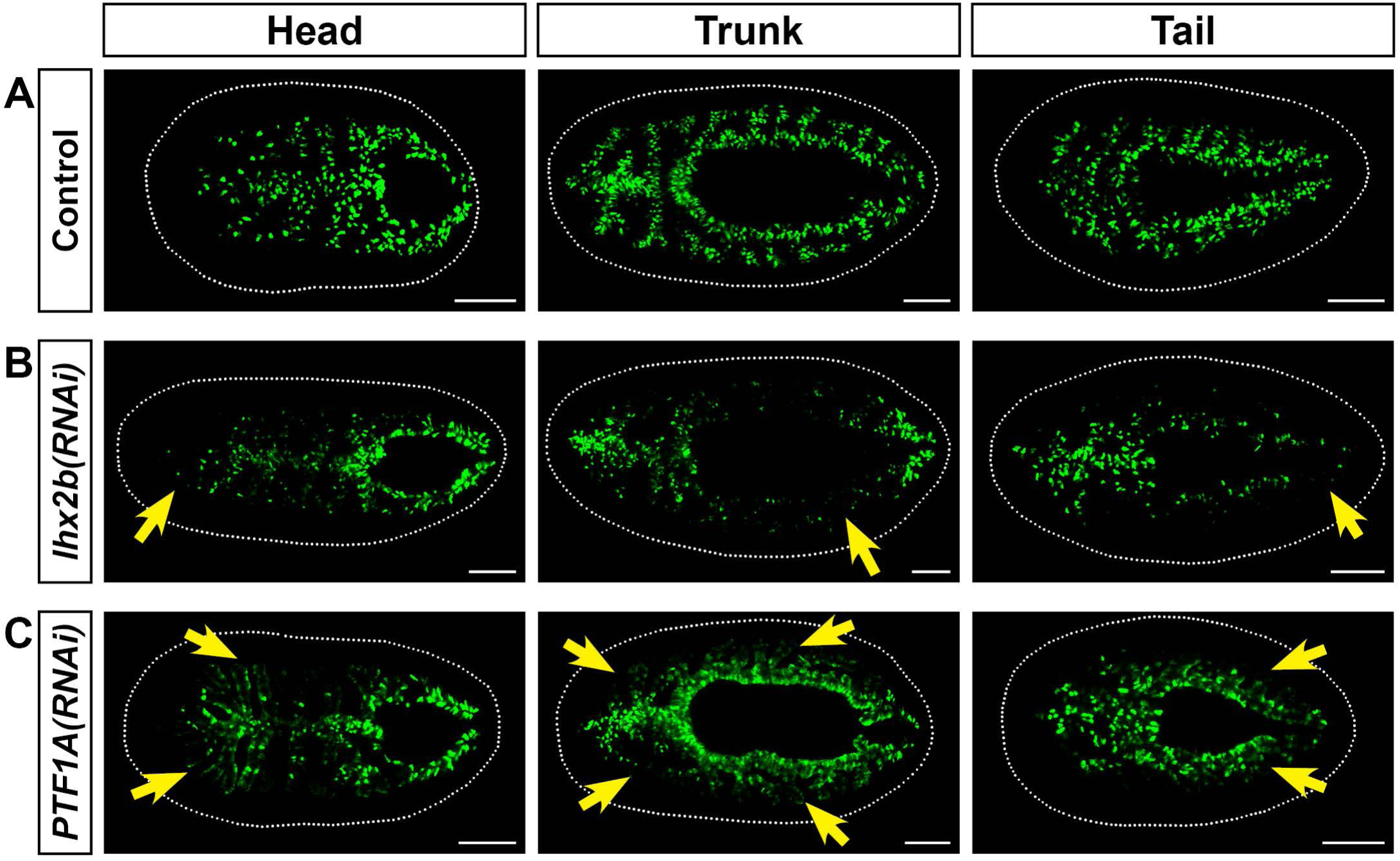
Additional genes whose knockdown affects goblet cell numbers in 7-day regenerates. **(A)** Control RNAi regenerates with normal numbers of goblet cells. **(B)** *Lhx2b* RNAi reduces numbers of goblet cells in pre-existing tissue, but not in the blastema. **(C)** *PTF1A* RNAi mildly reduces levels of lateral goblet cells. All in situ hybridizations to detect *npc2-*positive goblet cells (green). Detailed gene ID information is available in Table S1. Scale bars, 200 μm.

**Figure S11.**
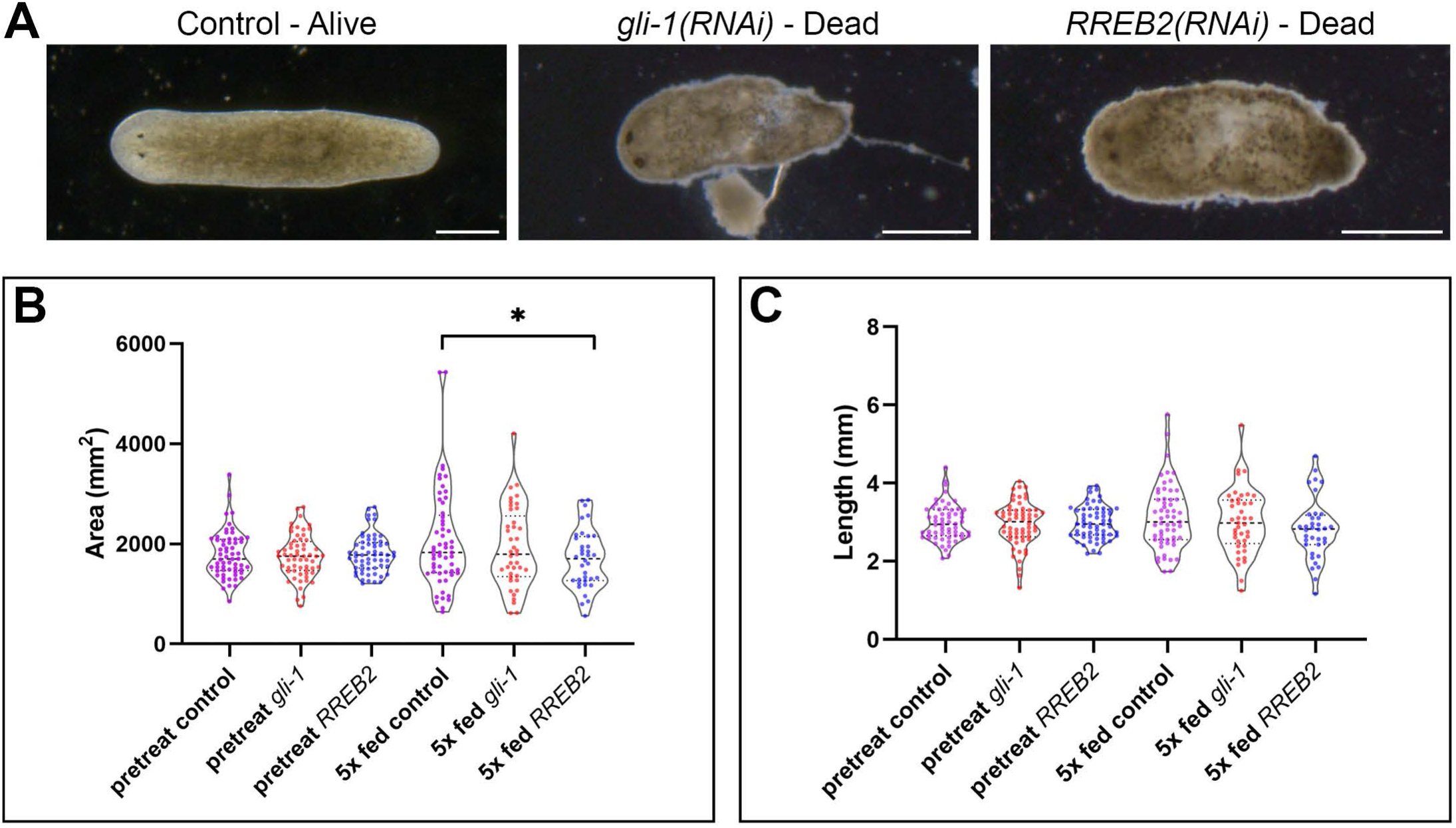
Phenotypes, area, and length of *gli-1* and *RREB2* knockdowns. **(A)** Example images of viability phenotypes observed during dsRNA feeding. **(B)** Area measurements immediately before the first dsRNA feeding and seven days after the fifth dsRNA feedings (one feeding per week). No significant differences were observed between pre-treatment area sizes (n=60 each), and no difference is observed between 5x fed control (n=57) and 5x fed *gli-1* (n=41, p-val = 0.4174), but a small difference is observed between 5x fed control and 5x fed *RREB2* (n=36, p-val = 0.0258). Non-eating animals and dead animals were not included in size analysis. **(C)** Length measurements from the same animals in **(B)** before and after dsRNA feeding. No significant differences between the control and the experiments were observed (5x fed control vs 5x fed *gli-1* p-value = 0.5010, 5x fed control vs. 5x fed *RREB2* p-value = 0.0769). Thick dashed line indicates the median, the thin dashed lines indicate the upper and lower quartiles.

**Figure S12.**
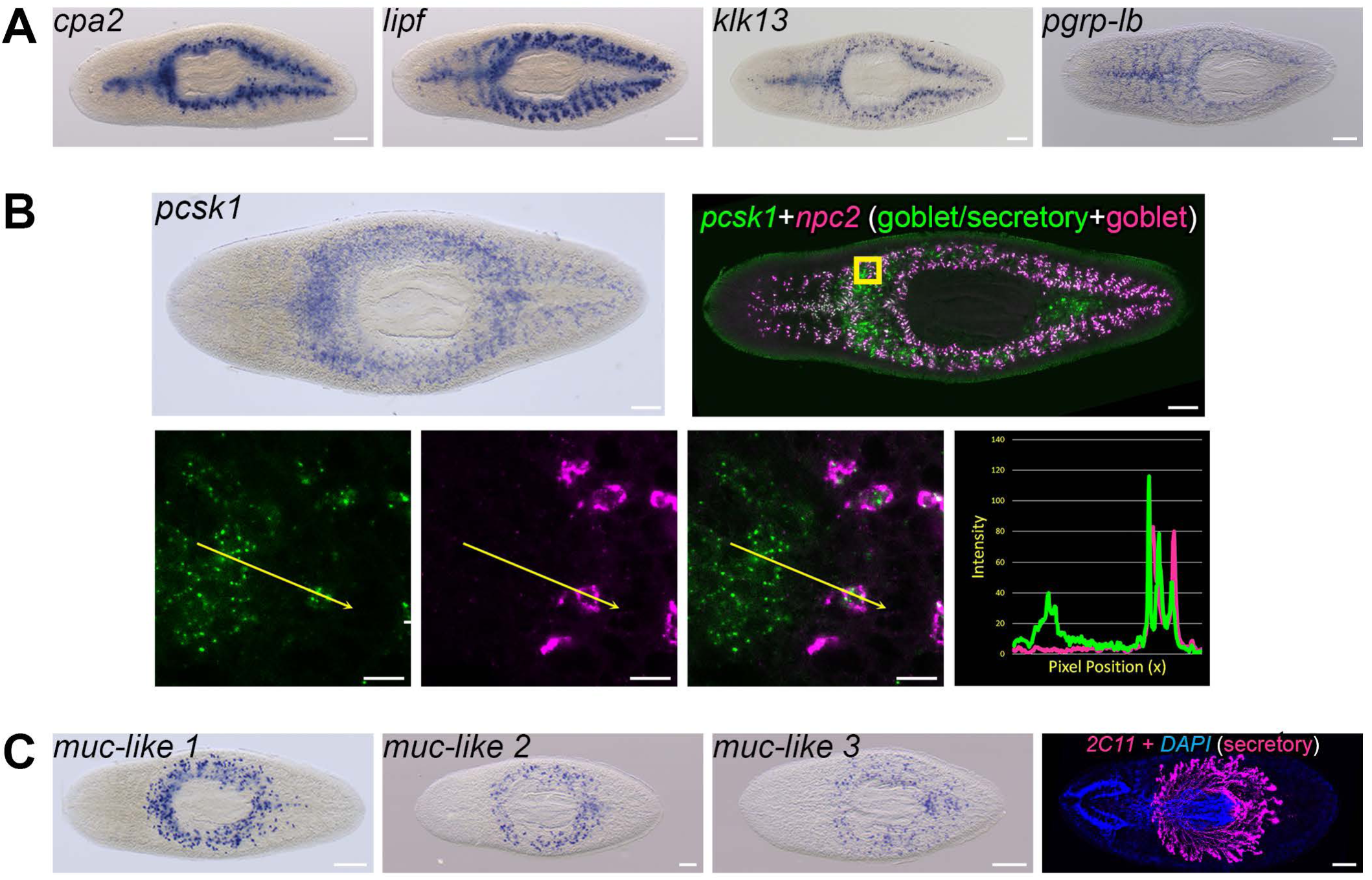
Gene expression by goblet cells suggests multiple physiological roles. **(A)** Whole-mount in situ hybridizations for *carboxypeptidase A2 (cpa2), gastric triacylglycerol lipase (lipf), kallikrein-13* (*klk13),* and *peptidoglycan recognition protein LB (pgrp-lb)*. **(B)** Neuroendocrine convertase 1 (*pcsk1*) mRNA expression. WISH, top left. dFISH, top right and magnified views (yellow box). *pcsk1* (green) co-expression with the pan-goblet marker *npc2* (magenta) demonstrates *pcsk1* is expressed in goblet cells as well as peripharyngeal secretory cells. **(C)** Expression of three *mucin*-like genes in secretory cells surrounding the pharynx (*muc-like-1/*dd_Smed_v6_17988_0_1, *muc-like-2/*dd_18786_0_1, and *muc-like-3/*dd_21309_0_1), and mAb 2C11 labeling peripharyngeal secretory cells and their projections into the pharynx (magenta) (bottom row). Detailed gene ID information is available in Table S1. Scale bars: whole animals 200 μm; magnified images (B), 10 µm.

**Figure S13.**
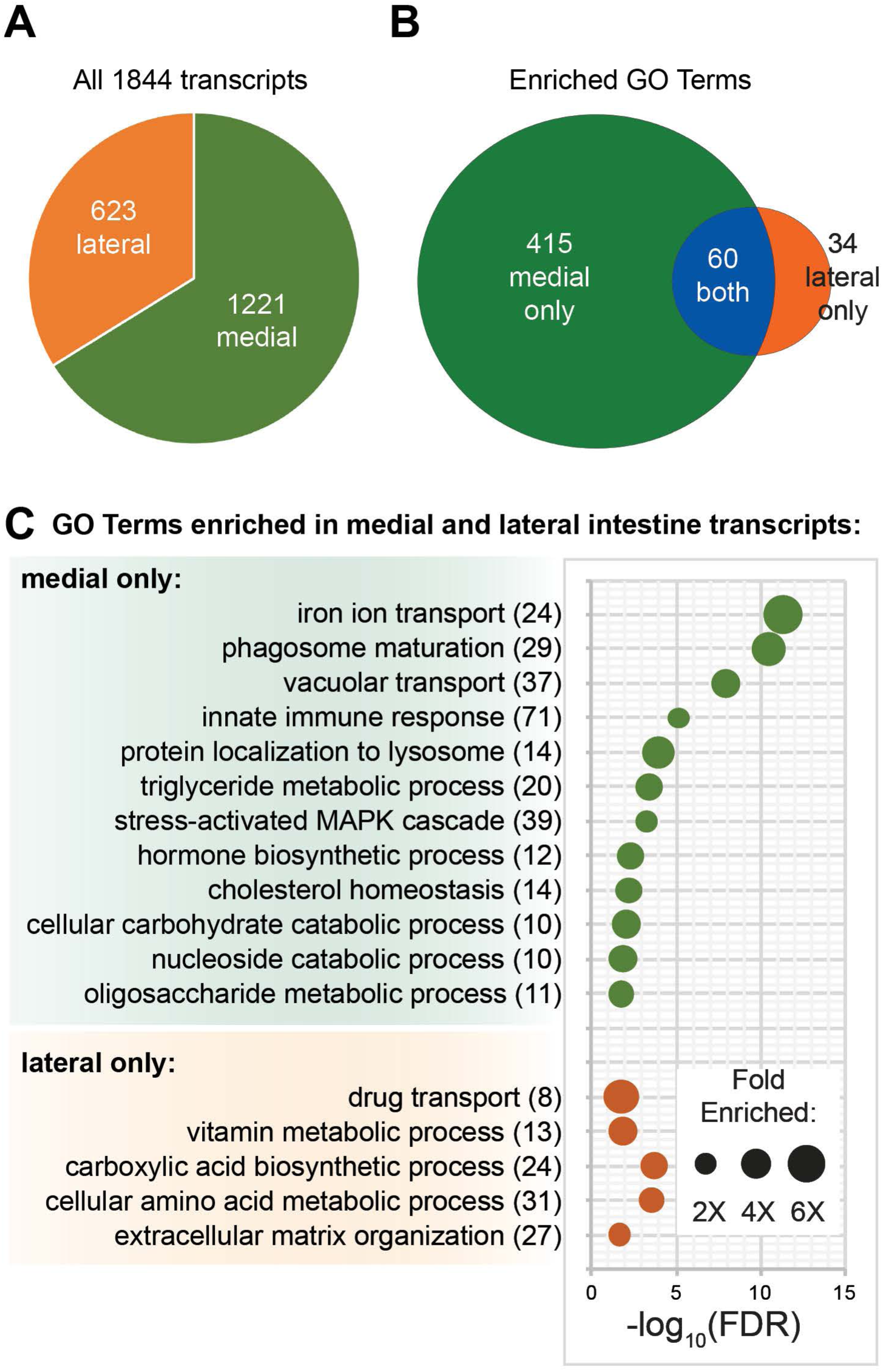
Biological Process GO terms enriched among medially and laterally enriched transcripts. **(A)** Pie chart showing the ratio of transcripts with higher fold changes in medial and lateral intestinal tissue. **(B)** Venn diagram showing the number of Biological Process GO terms enriched in medial, lateral, or both groups of transcripts. **(C)** Examples of GO terms over-represented in medial and lateral transcripts.

Table S1. Transcripts enriched in the planarian intestine.

Table S2. Gene ontology biological process term enrichment for intestinal transcripts.

Table S3. Intestine-enriched transcripts with potential roles in innate immunity.

Table S4. Planarian transcripts with reciprocal best hit orthologs enriched in human tissues.

Table S5. Comparison of transcripts enriched in laser-captured intestine and sorted intestinal phagocytes.

Table S6. RNA interference results for intestine-enriched transcription factors and mediolaterally enriched transcripts.

